# Renal mechanotransduction is an essential regulator of renin

**DOI:** 10.1101/2023.11.04.565646

**Authors:** Rose Z. Hill, Sepenta Shirvan, Sebastian Burquez, Adrienne E. Dubin, M. Rocio Servin-Vences, Jeffrey H. Miner, Ardem Patapoutian

## Abstract

The kidneys tightly control the composition of our internal environment to maintain homeostasis in the face of external variability. The regulation of blood volume begins in the kidneys and is essential for vertebrate life in terrestrial environments where salt and water availability are unpredictable^1,2^. Renin synthesis and release by juxtaglomerular granular cells of the kidney is the rate-limiting step in a hormonal cascade that modulates blood volume, filtration, and salt balance^2^. Renin is stimulated during hypovolemia and salt deprivation in response to chemical cues released from sympathetic efferent neurons and the macula densa onto the juxtaglomerular granular cells. Renin levels are also proposed to be modulated by mechanical forces elicited by changes in blood volume and/or pressure exerted upon juxtaglomerular cells^2–4^. However, the identity and significance of the juxtaglomerular mechanotransducer(s) was unknown. We found that the force-gated ion channel PIEZO2 is expressed in juxtaglomerular granular cells and in neighboring mesangial cells. Selective genetic ablation of PIEZO2 dysregulated the renin-angiotensin-aldosterone system by elevating renin, raising systemic blood pressure, inducing glomerular hyperfiltration, and exaggerating the hormonal response to volume depletion. These findings demonstrate that PIEZO2 contributes to renal blood volume sensing and kidney function *in vivo*.

## Main

The scientist and author Homer Smith wrote, “Our kidneys constitute the major foundation of our physiological freedom”^1^. Vertebrate animals control their blood volume through intersecting feedback loops spanning the renal, nervous, and cardiovascular systems^2,3,5^. These processes govern blood pressure, cardiovascular function, electrolyte balance, thirst, and fluid excretion through diverse mechanisms. Importantly, they permit organisms to adapt to physiologically demanding environments. For example, it is known that neuronal baroreceptors innervating the blood vessels detect mechanical forces exerted by pressure changes via mechanically gated PIEZO ion channels. This baroreceptor rapidly alters cardiac output, heart rate, and vascular resistance via engagement of the autonomic nervous system^3,5,6^. A non-neuronal baroreceptor in the kidney responds to volume depletion caused by salt scarcity, dehydration, or a substantial blood loss through activation of the renin-angiotensin-aldosterone system (RAAS), a hormonal cascade that increases vascular resistance and conserves electrolytes and water^2,7^. Renin is synthesized by and released from the juxtaglomerular granular (JG) cells, specialized vascular smooth muscle cells that decorate the afferent arteriole feeding the glomerulus. Renin production and release from JG cells are the rate-limiting steps of RAAS. Classical physiological studies showed that renin production and/or release into the circulation can be stimulated from JG cells by way of three main routes: 1) sympathetic efferent-derived norepinephrine released onto JG cells in response to neuronal baroreceptor activation^8,9^, 2) a reduction in tubular salt concentration that triggers neighboring cells of the macula densa to secrete prostaglandins onto the JG cells^2,10–13^, and, 3) mechanotransduction within JG cells from loss of afferent arteriolar stretch that stimulates renin release in an inversely calcium-dependent manner^2,14,15^.

Of these several means to increase plasma renin levels, the least understood is mechanosensing by the JG cells. A number of ion channels and pathways have been proposed to serve this role in JG cells^16–18^. However, the physiological role(s) of mechanically activated ion channels in the renal baroreceptor have not been rigorously tested *in vivo*. It is thought that stretching of the afferent arteriole mechanically stimulates the JG cells, driving calcium influx and suppressing renin, whereas volume depletion relaxes the arteriole and prevents calcium influx to stimulate renin^15^. Furthermore, while it has been hypothesized that the JG cells themselves are the mechanosensors^4,14^, they may be electrically coupled by gap junctions to neighboring mesangial cells^19–22^. As such, the cellular site of mechanotransduction cannot be assumed. Additionally, the *in vivo* consequences of loss of JG mechanosensitivity are unknown, particularly in light of parallel mechanisms to stimulate RAAS. In our study, we sought to elucidate the molecular identity and physiological significance of mechanotransduction within the JG apparatus (JGA).

### PIEZO2 is expressed in juxtaglomerular and mesangial cells of the kidney

Given the link between intracellular calcium levels and renin, we hypothesized that mechanically activated nonselective cation channels might underlie the intrinsically mechanosensitive component of the renal baroreceptor. PIEZO1 and PIEZO2 comprise a family of ion channels that are exclusively gated by membrane stretch, and are both necessary and sufficient for cellular responses to a range of physiologically salient mechanical stimuli^23–25^. PIEZO1 is endogenously expressed in many tissue types including vascular endothelium and smooth muscle, and is important for vascular development and function^26,27^. PIEZO2 is mainly expressed in peripheral sensory neurons and specialized accessory sensory cells and is required for gentle touch sensation, proprioception, and gut and bladder function^28–31^. Together, PIEZO1 and PIEZO2 in vagal sensory neurons initiate the neuronal baroreflex through sensing of blood pressure in the aortic arch^5^. To establish the expression patterns of PIEZOs in the mouse kidney, we first examined the localization of mRNA transcripts of mechanosensory PIEZO channels using single molecule fluorescence in situ hybridization (smFISH). We found that *Piezo2* and not *Piezo1* was expressed in *Ren1*-expressing JG cells of the JGA and in *Pdgfrb*-expressing mesangial cells of the glomerulus^32,33^ (Fig. 1a-c). Using a *Piezo2^GFP-Cre^* fusion knock-in mouse^29^, we observed a pattern of fusion protein expression restricted to glomerular cells (Fig. 1d) as well as robust glomerular and JG cell labeling when crossed with the *Ai9^fl/fl^*tdTomato reporter line^34^ (Fig. 1e). By contrast, immunohistochemistry (IHC) of PIEZO1-tdTomato C-terminal fusion protein from a *Piezo1^tdTomato^* knock-in mouse line showed expression largely restricted to basal aspects of a subset of tubular epithelial cells as previously reported^35^ (Extended Data Fig. 1a). Our findings are in agreement with single-nucleus RNA-seq databases of mouse and human kidney tissues that support the expression of *Piezo2* and not *Piezo1* in mesangial and renin-expressing JG cells^36^. To further validate those results, we performed RNAscope for *PIEZO2* and *PDGFRB* on kidney sections from a healthy human donor. We observed overlap of the two transcripts’ expression patterns in kidney cortex (Extended Data Fig. 1b).

**Figure 1.**
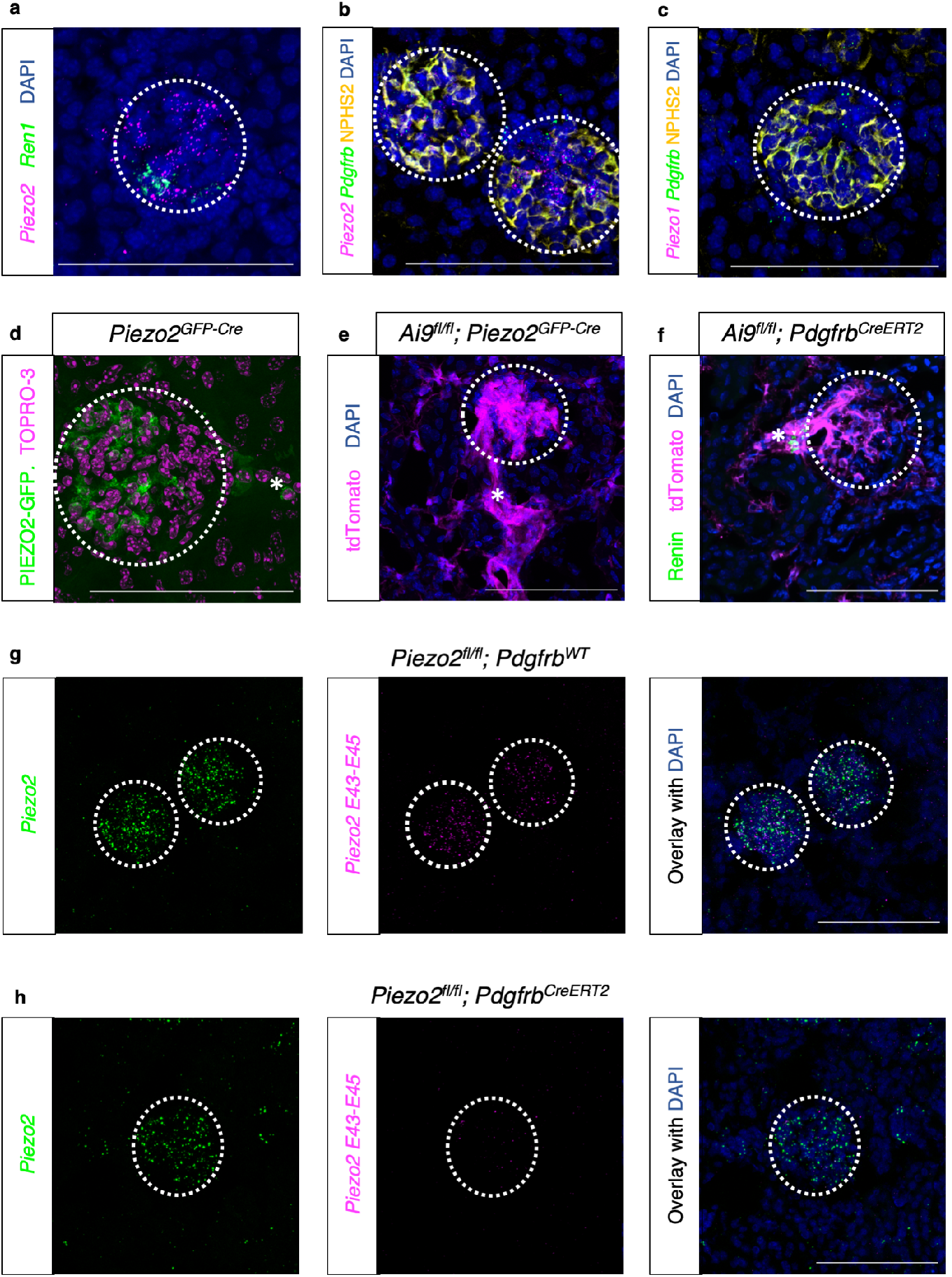
PIEZO2 is expressed in JG and mesangial cells of the kidney. **a**, smFISH of sectioned mouse kidney for *Piezo2*, *Ren1*, and counterstained with DAPI. **b**, smFISH with IHC of sectioned mouse kidney for *Piezo2*, *Pdgfrb*, anti-NPHS2, and counterstained with DAPI. **c**, smFISH with IHC of sectioned mouse kidney for *Piezo1*, *Pdgfrb*, anti-NPHS2, and counterstained with DAPI. **d**, Sectioned mouse kidney stained with anti-GFP antibody and counterstained with TO-PRO-3. **e**, Sectioned mouse kidney with native tdTomato fluorescence and counterstained with DAPI. **f**, Sectioned mouse kidney with native tdTomato fluorescence, stained with anti-Renin antibody, and counterstained with DAPI. Asterisk (*) indicates extraglomerular expression at putative Renin+ vascular pole. **g-h**, smFISH of sectioned mouse kidney for *Piezo2* (left), floxed exon-specific probe *Piezo2 E43-45* (center), and merged image counterstained with DAPI (right). Scale bars = 100 µm. Dotted circles indicate renal corpuscles. Each experiment was repeated on at least N=2 mice.

In order to target PIEZO2 for genetic ablation in the mouse kidney, we turned to an inducible pericyte Cre recombinase mouse line, *Pdgfrb^CreERT2^*, that selectively targets specialized vascular smooth muscle cells that include mesangial and renin-expressing JG cells^37^ after tamoxifen administration to adult mice to induce recombination (Fig. 1f, Extended Data Fig. 1c-d). Indeed, we found that after crossing this line with *Piezo2^fl/fl^*mice, tamoxifen ablated *Piezo2* expression in the kidney of *Pdgfrb^CreERT2^*but not *Pdgfrb^WT^* animals, confirmed using an smFISH assay with probes against either the entire transcript (*Piezo2*) or the *loxp*-flanked exons (*Piezo2 E43-E45*, Fig. 1g-h). We observed additional patterns of Cre recombination in *Ren^Cre^* and *FoxD1^GFP-Cre^* mouse lines after crossing with the *Ai9^fl/fl^* tdTomato reporter strain. In line with a published study, *Ren^Cre^* targeted renin-expressing putative JG cells of the JGA and renin-negative cells of the renin lineage along the glomerular arterioles of the kidney^38^, while largely sparing mesangial cells (Extended Data Fig. 1e-f). We observed that the stromal progenitor Cre line *FoxD1^GFP-Cre^*targeted a broad population encompassing mesangial cells, renin-expressing cells, and fibroblasts, while sparing tubules and endothelial cells as previously reported^39^ (Extended Data Fig. 1g-h). While our initial testing of a conditional *Ren^CreER^*reporter line used previously for lineage tracing^40^ showed selective targeting of adult renin-expressing cells, we observed only partial recombination in these cells after tamoxifen administration, rendering it unsuitable for genetic loss-of-function studies *in vivo* (Extended Data Fig. 1i). It is worth noting that while none of our Cre lines (*Pdgfrb^CreERT2^*, *Ren^Cre^*, and *FoxD1^GFP-Cre^*) on their own are selective for renin-expressing JG cells, PIEZO2, unlike PIEZO1, is expressed in only handful of cell types in the body and mainly found within the peripheral nervous system^5,28,41^. Although PIEZO2 expression has been sporadically observed in vascular endothelial cells^42^ (its function in these cells is unknown), none of our Cre lines target this particular cell type and we were unable detect *Piezo2* transcript expression in glomerular capillary endothelial cells. As such, by using *Pdgfrb^CreERT2^*, *Ren^Cre^*, and *FoxD1^GFP-Cre^*in combination we can infer the function of PIEZO2 in JG cells, which are uniquely targeted by all three of the genetic strategies.

### Renal PIEZO2 is an essential regulator of plasma renin

We next sought to investigate the consequences of loss of functional PIEZO2 to renin levels and RAAS. Since the primary role of the JG cells is to synthesize and release renin, we measured renin in plasma harvested from naïve mice. Initially, to avoid potential confounds due to developmental phenotypes that could be present in *Ren^Cre^* or *FoxD1^GFP-Cre^*conditional mutant mice, and to target all PIEZO2-expressing renal cells (both mesangial and putative JG cells), we examined the inducible *Piezo2^fl/fl^; Pdgfrb^CreERT2^* conditional knockout mice. We observed a significant increase in plasma renin levels in the *Piezo2^fl/fl^; Pdgfrb^CreERT2^* mice compared to *Piezo2^fl/fl^; Pdgfrb^WT^*wild-type littermate controls that also received tamoxifen (Fig. 2a). This effect was not exacerbated in *Piezo1^fl/fl^; Piezo2^fl/fl^; Pdgfrb^CreERT2^* mice lacking both ion channels (Extended Data Fig. 2a), consistent with our earlier observation that PIEZO2 and not PIEZO1 is expressed in mesangial and JG cells. The elevated renin levels were phenocopied in *Piezo2^fl/fl^; FoxD1^GFP-Cre^*animals, suggesting that loss of PIEZO2 in stromal cells was sufficient to reproduce the phenotype and was not likely due to PIEZO2 in non-stromal populations including tubular and endothelial cells (Extended Data Fig. 2b). Notably, we did not observe elevations in downstream RAAS hormones aldosterone and angiotensin II (Fig. 2b-c) in *Piezo2^fl/fl^; Pdgfrb^CreERT2^* mice, suggesting that the elevated renin levels were insufficient to trigger induction of downstream RAAS hormones under baseline conditions. While hyperreninemia is sometimes associated with hypertension, this is typically attributed to a systemic elevation in angiotensin II^43,44^, which was not observed in plasma from the conditional knockout mice (Fig. 2c). We measured the systemic blood pressure *Piezo2^fl/fl^; Pdgfrb^WT^* and *Pdgfrb^CreERT2^*mice using the radiotelemetry-validated volume pressure recording (VPR) method^45^ and found a significant elevation in *Pdgfrb^CreERT2^* mice that was not indicative of hypertension (Fig. 2d, Extended Data Figs. 3a-b, 4a-c). The effect was less than that reported in transgenic rodent models with elevated RAAS signaling^44^, and may reflect the lack of significantly elevated angiotensin II in the naïve PIEZO2 conditional knockout mice. However, it is possible that the elevated baseline renin observed in the conditional knockout mice contributes to sporadic or mild elevations in angiotensin II not captured by our enzyme-based measurement methods, but which are still capable of triggering mildly elevated blood pressure.

**Fig. 2.**
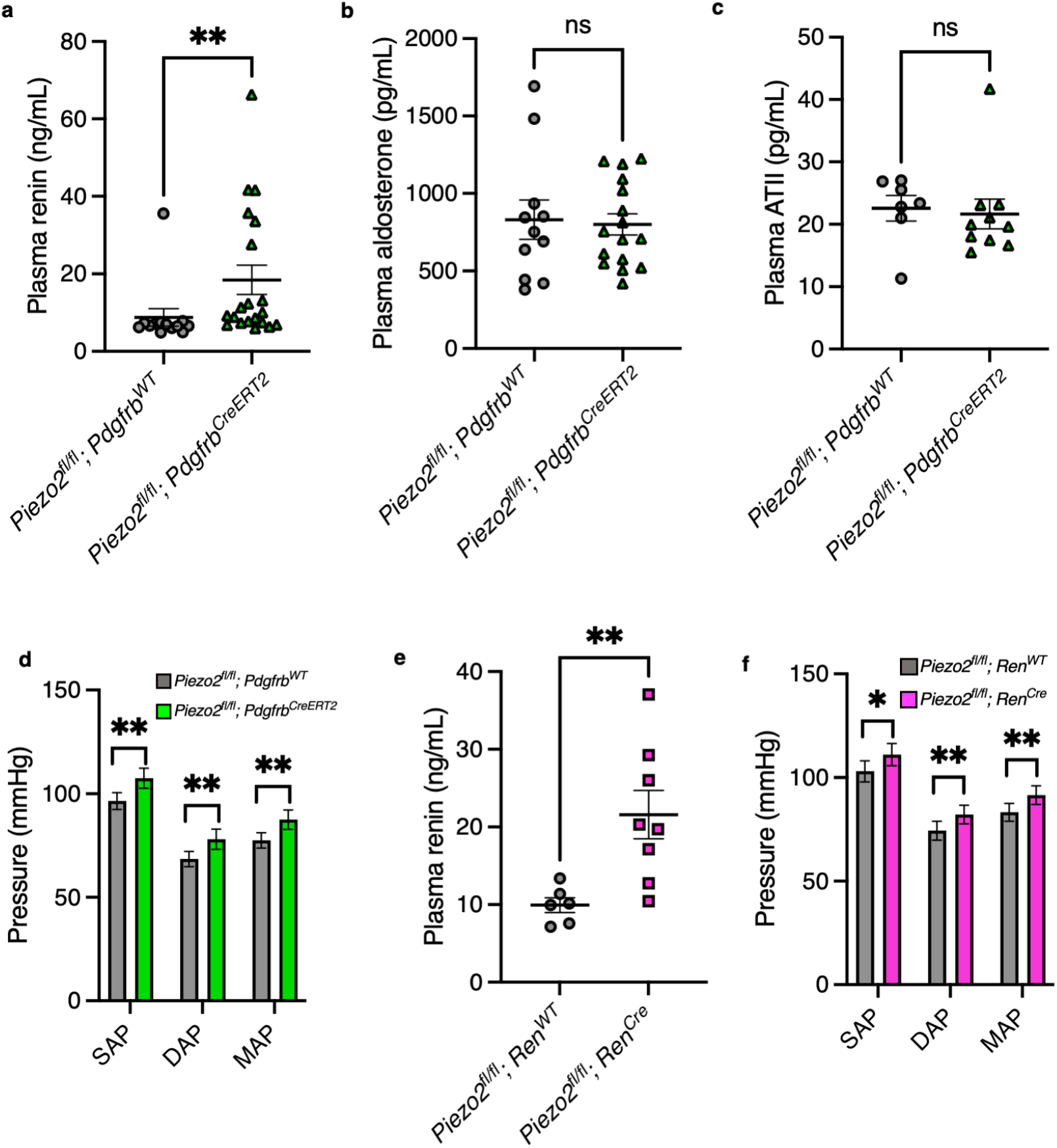
Renal PIEZO2 is an essential regulator of plasma renin. **a**, Plasma renin levels in *Piezo2^fl/fl^; Pdgfrb^WT^* versus *Piezo2^fl/fl^; Pdgfrb^CreERT2^* animals (Mann–Whitney: \*\**p* = 0.0019, U = 48; n = 20 *Pdgfrb^WT^* and 13 *Pdgfrb^CreERT2^* mice). **b**, Plasma aldosterone levels in *Piezo2^fl/fl^; Pdgfrb^WT^* versus *Piezo2^fl/fl^; Pdgfrb^CreERT2^* animals (Mann–Whitney: *p* = 0.8653, U = 84; n = 11 *Pdgfrb^WT^* and 16 *Pdgfrb^CreERT2^* mice). **c**, Plasma angiotensin II (ATII) levels in *Piezo2^fl/fl^; Pdgfrb^WT^* versus *Piezo2^fl/fl^; Pdgfrb^CreERT2^* animals (Mann–Whitney: *p* = 0.1932, U = 21; n = 7 *Pdgfrb^WT^* and 10 *Pdgfrb^CreERT2^* mice). **d**, Systemic blood pressure (systolic/SAP; diastolic/DAP; and mean arterial pressure/MAP) measured using the VPR system in *Piezo2^fl/fl^; Pdgfrb^WT^* versus *Piezo2^fl/fl^; Pdgfrb^CreERT2^* animals (two-tailed nested *t*-tests (left to right): \*\**p_SAP_* = 0.0013, *t* = 3.899, d.f. = 16; \*\**p_DAP_* = 0.0056, *t* = 3.197, d.f. = 16; \*\**p_MAP_* = 0.0027, *t* = 3.546, d.f. = 16; n = 10 *Pdgfrb^WT^* and 8 *Pdgfrb^CreERT2^* mice). **e**, Plasma renin levels in *Piezo2^fl/fl^; Ren^WT^* versus *Piezo2^fl/fl^; Ren^Cre^* animals (Mann–Whitney: \*\**p* = 0.0047, U = 3; n = 6 *Ren^WT^* and 8 *Ren^Cre^* mice). **f**, Systemic blood pressure measured in *Piezo2^fl/fl^; Ren^WT^* versus *Piezo2^fl/fl^; Ren^Cre^* animals (two-tailed nested *t*-tests (left to right): \**p_SAP_* = 0.0147, *t* = 2.716, d.f. = 17; \*\**p_DAP_* = 0.0097, *t* = 2.913, d.f. = 17; \*\**p_MAP_* = 0.0060, *t* = 3.137, d.f. = 17; n = 9 *Ren^WT^* and 10 *Ren^Cre^* mice). Each experiment was performed on at least two independent cohorts of mice, and error bars represent mean ± s.e.m.

To determine whether mesangial and/or renin-lineage cells contribute to the elevated renin in PIEZO2 conditional mutants, we measured plasma renin and blood pressure in *Piezo2^fl/fl^; Ren^Cre^*conditional knockouts and littermate controls. Our findings in this strain phenocopied those observed with *Pdgfrb^CreERT2^*-mediated deletion of PIEZO2 (Figs. 2e-f, Extended Data Figs. 5a-b, 6a-c). As *Ren^Cre^* targets JG cells and other cells of renin lineage within the kidney but spares the majority of mesangial cells^38,40^, we conclude from these experiments that the phenotype is primarily driven by PIEZO2 in renin+ JG cells rather than in mesangial cells, which are known to play key roles in the pathogenesis of glomerular disease and are relatively unstudied in the context of RAAS^46^, though we cannot completely rule out their involvement.

### Renin regulates the GFR via Ang(1-7)/Mas in a PIEZO2-dependent manner

Another important function of renin is modulating the glomerular filtration rate (GFR) in response to changes in renal blood flow and macula densa signaling^4,47,48^. We observed that *Piezo2^fl/fl^; Pdgfrb^CreERT2^* mice exhibited glomerular hyperfiltration in an assay in which FITC-sinistrin (a freely filterable molecule that is neither reabsorbed into the blood nor secreted through the peritublar capillaries) fluorescence signal decay is observed by continuous transdermal measurement^49^ (Fig. 3a-b), consistent with a dysregulation of renin. The GFR values of the conditional knockout mice were similar to those observed in genetic models of hyperfiltration^50^. Whereas kidney disease is typically associated with a low GFR, early in its pathogenesis certain diseases such as diabetes trigger glomerular hyperfiltration^51,52^. As such, we also explored the possibility that the *Piezo2^fl/fl^; Pdgfrb^CreERT2^* mice had renal disease by re-measuring GFR in the same adult mice eight months after the initial experiments. We found that the GFR of the *Piezo2^fl/fl^; Pdgfrb^CreERT2^* mice declined as expected with age^53–55^, comparable to the decline observed in wild-type littermates (Extended Data Fig. 7a). The conditional knockout mice maintained elevated GFR compared to controls, suggesting that the observed phenotype could be due to elevated renin rather than kidney disease (Extended Data Fig. 7a-b). Additionally, the conditional knockout mice did not exhibit elevated urinary albumin, a hallmark of kidney glomerular disease (Extended Data Fig. 7c). We also observed that the elevated GFR was phenocopied in *Piezo2^fl/fl^; Ren^Cre^* conditional knockouts, suggesting the phenotype was mediated by renin-expressing cells rather than glomerular mesangial cells (Fig. 3c). Importantly, loss of PIEZO1 and PIEZO2 in peripheral neuronal baroreceptors using *Piezo1^fl/fl^; Piezo2^fl/fl^; SNS^Cre^* ^56^ did not induce a GFR phenotype (Extended Data Fig. 7d), suggesting that these findings are independent of any indirect effects of neural baroreception on kidney function^3,5,57^, for example, due to modulation of sympathetic norepinephrine release onto the JGA. Examination of PAS- and H&E-stained sections showed that neither *Piezo2^fl/fl^; Pdgfrb^CreERT2^* nor *Piezo2^fl/fl^; Ren^Cre^* kidneys exhibited consistent histological abnormalities compared to controls that were indicative of glomerular disease or dysfunction (Extended Data Fig. 7e-h), further supporting the idea that the elevated GFR was not due to overt glomerular disease.

**Fig. 3.**
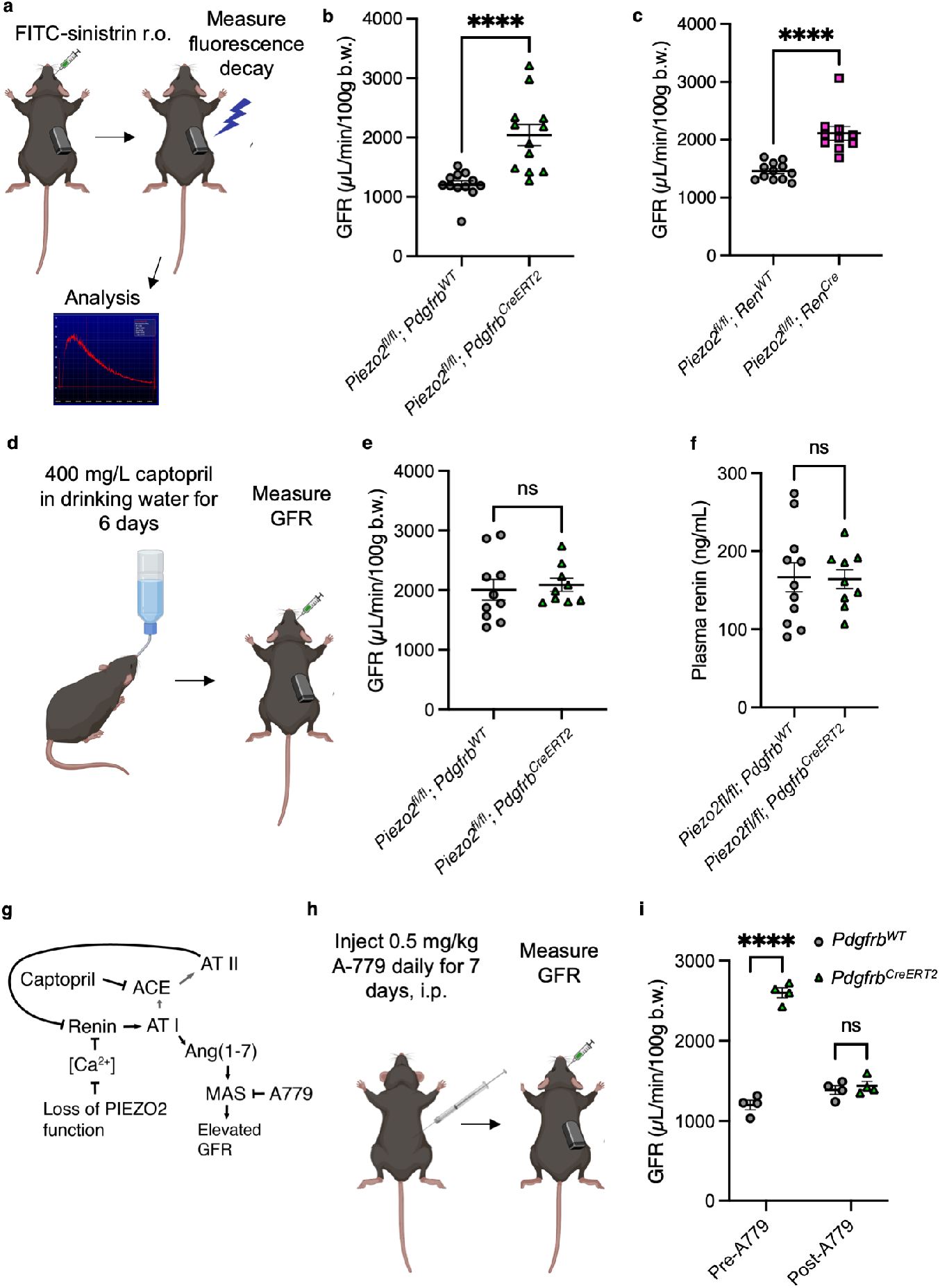
Renin regulates the GFR via Ang(1-7)/Mas in a PIEZO2-dependent manner. **a**, FITC-Sinistrin transdermal GFR measurement in mice. **b**, GFR in *Piezo2^fl/fl^; Pdgfrb^WT^* versus *Piezo2^fl/fl^; Pdgfrb^CreERT2^* animals (Mann–Whitney: \*\*\*\**p* < 0.0001, U = 7; n = 12 *Pdgfrb^WT^* and 12 *Pdgfrb^CreERT2^* mice). **c**, GFR in *Piezo2^fl/fl^; Ren^WT^* versus *Piezo2^fl/fl^; Ren^Cre^* animals (Mann–Whitney: \*\*\*\**p* < 0.0001, U = 1; n = 12 *Ren^WT^* and 10 *Ren^Cre^* mice). **d**, Captopril experiment. **e**, GFR after daily administration of 400 mg/L captopril in the drinking water of *Piezo2^fl/fl^; Pdgfrb^WT^* and *Piezo2^fl/fl^; Pdgfrb^CreERT2^* animals for six days (Mann–Whitney: *p* = 0.4470, U = 35; n = 10 *Pdgfrb^WT^* and 9 *Pdgfrb^CreERT2^* mice). **f**, Plasma renin levels in *Piezo2^fl/fl^; Pdgfrb^WT^* versus *Piezo2^fl/fl^; Pdgfrb^CreERT2^* animals after seven days of captopril (Mann–Whitney: *p* = 0.8820, U = 47; n = 11 *Pdgfrb^WT^* and 9 *Pdgfrb^CreERT2^* mice). **g**, Schematic of the role of renin in GFR. **h**, A779 Mas blockade experiment. **i**, GFR before (pre-A779) and after (post-A779) treatment with A779 in *Piezo2^fl/fl^; Pdgfrb^WT^* versus *Piezo2^fl/fl^; Pdgfrb^CreERT2^* animals (two-way ANOVA: \*\*\*\**p_interaction_* < 0.0001, F(1,6) = 248.9; Sidak’s multiple comparisons: \*\*\*\**p_pre-A779_* < 0.0001, *p_post-A779_* = 0.7724; n = 4 *Pdgfrb^WT^* and 4 *Pdgfrb^CreERT2^* mice). Each experiment was performed on at least two independent cohorts of mice, and error bars represent mean ± s.e.m.

We postulated that the elevated GFR, as it did not arise from overt kidney disease nor from loss of PIEZO2 in mesangial cells, could be caused by aberrant signaling downstream of renin. Elevated renin is a consequence of administration of captopril, an angiotensin converting enzyme (ACE) inhibitor commonly prescribed to treat hypertension that disrupts feedback inhibition of angiotensin II on renin production^58^. The effects of this drug on GFR are unclear^59–62^. We reasoned that if captopril causes elevated plasma renin, then we would expect captopril to enhance the GFR of wild-type mice to a level comparable to that of the conditional knockouts (Fig. 2a). Indeed, we observed that six days of captopril administration (Fig. 3d) elevated the GFR of wild-type mice such that it was not significantly different from the *Piezo2^fl/fl^; Pdgfrb^CreERT2^* littermates also receiving captopril (Fig. 3e). Consistent with captopril’s known effects, we observed elevated plasma renin (Fig. 3f) and decreased aldosterone (Extended Data Fig. 8a) in mice of both genotypes on day seven^58^. It was of note that the renin levels observed in treated *Piezo2^fl/fl^; Pdgfrb^CreERT2^*animals were higher than those observed earlier in untreated animals (Fig. 2a), however, their GFR values were within a similar range (Figs. 3b,e). Taken together, our results suggest that elevation of plasma renin in the absence of elevated angiotensin II is sufficient to elevate the GFR. Our observation that the higher renin levels elicited by captopril in treated animals did not raise the GFR beyond that of untreated conditional mutant mice additionally suggests that there is a limit to renin’s ability to increase the GFR.

How does elevated renin increase the GFR? The relationship between RAAS activation and GFR is complex. For example, angiotensin II produced by RAAS activation can have opposing effects on GFR through differential effects on glomerular arteriole constriction and mesangial cell contractility ^61–64^. As our conditional knockout mice had normal angiotensin II levels, we did not suspect this molecule was playing a role in elevating the GFR of these animals. Notably, the related peptide Ang(1-7) is produced downstream of renin’s cleavage of angiotensinogen to angiotensin I and its subsequent conversion to Ang(1-7) by specialized proteases^65,66^ (Fig. 3g). Ang(1-7) acts via the G-protein coupled receptor (GPCR) Mas within in the glomerular vasculature to increase the GFR^67,68^. Intriguingly, this regulation by the Ang(1-7)/Mas axis is reported to occur under salt-depleted conditions^68^, as elevated renin drives Ang(1-7) production, and can occur after treatment with ACE inhibitors such as captopril that similarly drive elevated Ang(1-7) through loss of angiotensin I conversion to angiotensin II^66^ (Fig. 3g). We examined whether the elevated GFR could be a direct consequence of elevated renin and subsequent Ang(1-7)/Mas signaling in the *Piezo2^fl/fl^; Pdgfrb^CreERT2^* mice. We first tested whether Mas signaling via Ang(1-7) contributed to elevated GFR in the cKO mice. We measured the GFR before and after selectively blocking Mas signaling via daily administration of the pharmacological antagonist A779^67–70^ (Fig. 3h). Strikingly, Mas blockade fully rescued normal GFR in conditional knockout mice and had no effect on the GFR of littermate controls (Fig. 3i), demonstrating that Mas signaling underlies the elevation in GFR caused by loss of PIEZO2. Our findings with the captopril and A779 experiments establish a molecular pathway by which PIEZO2-dependent renin regulation is required for maintenance of a GFR within the normal range via Ang(1-7)/Mas signaling.

### PIEZO2 acts as a brake on the hormonal response to hypovolemia

A primary function of renin is to activate RAAS to conserve bodily salt and water during hypovolemia when blood volume is depleted. To test whether renal PIEZO2-dependent mechanotransduction plays a role in hypovolemia-evoked RAAS, we subjected *Piezo2^fl/fl^; Pdgfrb^CreERT2^*and littermate controls to the polyethylene glycol (PEG)-evoked hypovolemia model^71^ (Fig. 4a), where PEG draws bodily fluid into the subcutaneous space to cause a volume depletion from the blood into a secondary compartment without directly affecting salt balance, as with administration of the loop diuretic furosemide^8^. We found that the PEG-injected *Piezo2^fl/fl^; Pdgfrb^CreERT2^* mice exhibited an exaggerated hormonal response to hypovolemia, with elevated renin, angiotensin II, and aldosterone compared to littermate controls (Figs. 4b-d). The renin levels we measured were greater than that seen under normovolemic conditions (Fig. 2), and might explain the stronger induction of angiotensin II and aldosterone in conditional knockouts that were not seen in normovolemic *Piezo2^fl/fl^; Pdgfrb^CreERT2^* mice. Enhanced hypovolemia-evoked renin and aldosterone levels were phenocopied with the *Piezo2^fl/fl^; Ren^Cre^*and *FoxD1^Cre^* strains affecting the renin lineage and stromal progenitors (Figs. 4e-h), suggesting that loss of functional PIEZO2 in the renin-lineage alone was sufficient to drive the effect. Similar to our findings under naïve conditions, loss of both PIEZOs using *Piezo1^fl/fl^; Piezo2^fl/fl^; Pdgfrb^CreERT2^*had no further effect beyond loss of PIEZO2 alone (Extended Data Fig. 9a). No difference was observed between conditional knockout and controls with dual loss of PIEZO1 and PIEZO2 in peripheral neuronal baroreceptors (Extended Data Fig. 9b). We conclude from these experiments that PIEZO2 in renin-expressing cells blunts the RAAS response to hypovolemia.

**Figure 4.**
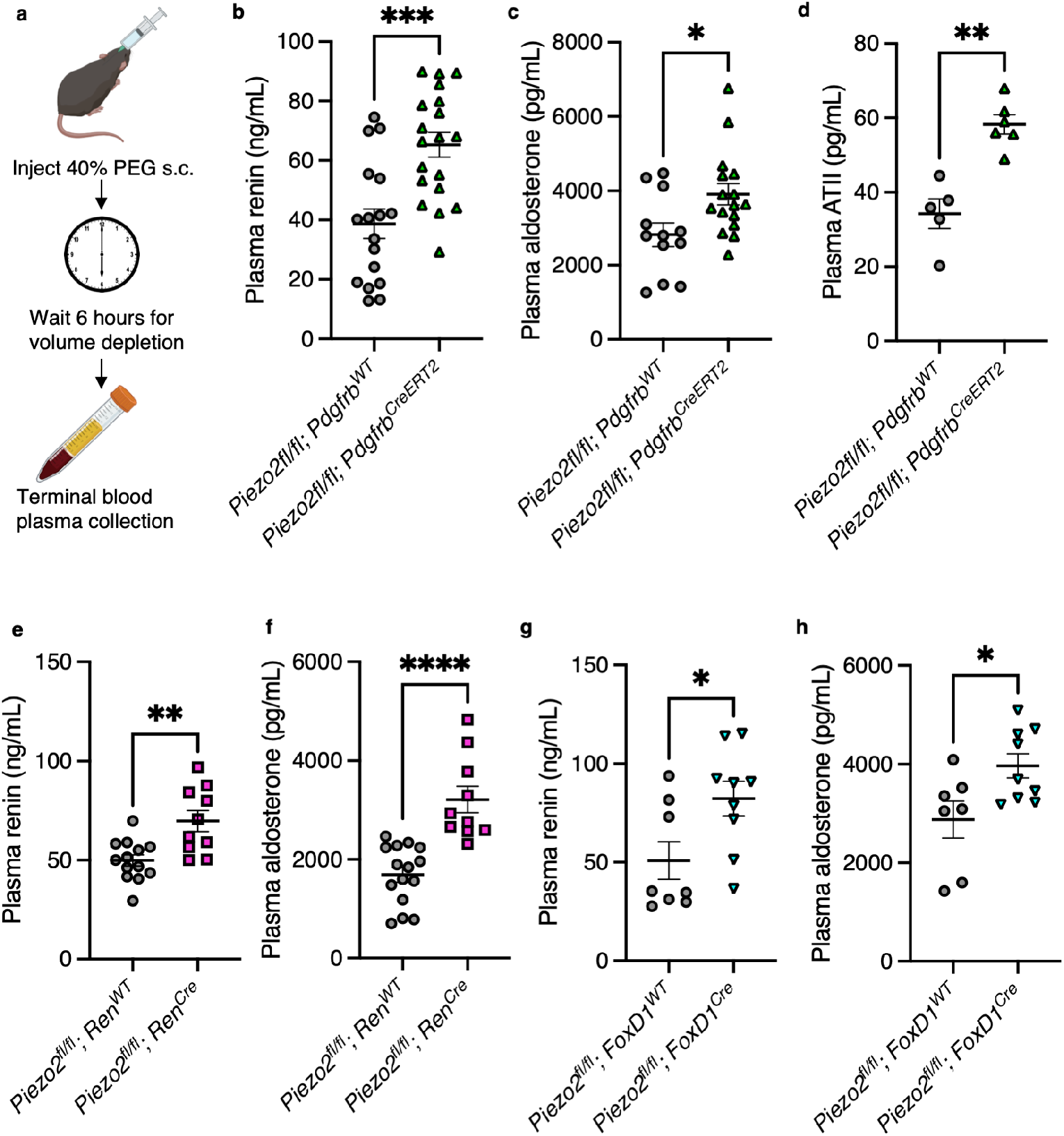
PIEZO2 in renin-expressing cells modulates the hormonal response to hypovolemia. **a**, The subcutaneous PEG model of hypovolemia. **b**, Plasma renin levels in *Piezo2^fl/fl^; Pdgfrb^WT^* versus *Piezo2^fl/fl^; Pdgfrb^CreERT2^* animals six hours following PEG (Mann– Whitney: \*\*\**p* = 0.0003, U = 53; n = 17 *Pdgfrb^WT^* and 19 *Pdgfrb^CreERT2^* mice). **c**, Plasma aldosterone levels in *Piezo2^fl/fl^; Pdgfrb^WT^* versus *Piezo2^fl/fl^; Pdgfrb^CreERT2^* animals six hours following PEG (Mann–Whitney: \**p* = 0.0257, U = 48; n = 12 *Pdgfrb^WT^* and 16 *Pdgfrb^CreERT2^* mice). **d**, Plasma ATII levels in *Piezo2^fl/fl^; Pdgfrb^WT^* versus *Piezo2^fl/fl^; Pdgfrb^CreERT2^* animals six hours following PEG (Mann–Whitney: \*\**p* = 0.0043, U = 0; n = 5 *Pdgfrb^WT^* and 6 *Pdgfrb^CreERT2^* mice). **e**, Plasma renin levels in *Piezo2^fl/fl^; Ren^WT^* versus *Piezo2^fl/fl^; Ren^Cre^* animals six hours following PEG (Mann–Whitney: \*\**p* = 0.0025, U =18; n = 13 *Ren^WT^* and 10 *Ren^Cre^* mice). **f**, Plasma aldosterone levels in *Piezo2^fl/fl^; Ren^WT^* versus *Piezo2^fl/fl^; Ren^Cre^* animals six hours following PEG (Mann– Whitney: \*\*\*\**p* < 0.0001, U =2; n = 15 *Ren^WT^* and 10 *Ren^Cre^* mice). **g**, Plasma renin levels in *Piezo2^fl/fl^; FoxD1^WT^* versus *Piezo2^fl/fl^; FoxD1^Cre^* animals six hours following PEG (Mann–Whitney: \**p* = 0.0360, U =14; n = 8 *FoxD1^WT^* and 9 *FoxD1^Cre^* mice). **h**, Plasma aldosterone levels in *Piezo2^fl/fl^; FoxD1^WT^* versus *Piezo2^fl/fl^; FoxD1^Cre^* animals six hours following PEG (Mann–Whitney: \**p* = 0.0418, U =12; n = 7 *FoxD1^WT^* and 9 *FoxD1^Cre^* mice). Each experiment was performed on at least two independent cohorts of mice, and error bars represent mean ± s.e.m.

### PIEZO2 contributes to RAAS independently of sympathetic and macula densa function

We next sought to determine how the contribution of PIEZO2 to RAAS is weighted against that of the other pathways regulating renin synthesis and release from JG cells, namely efferent sympathetic and macula densa signaling. To address this question, we designed an experiment in which we ablated or blocked these two respective arms through simultaneous chemical sympathectomy using the drug 6-hydroxydopamine^72,73^ (6-OHDA) and acute blockade of synthesis of prostaglandins, the primary macula densa-derived chemical signal stimulating renin, using the cyclooxygenase-1 and −2 inhibitor indomethacin^12,13^. After chemical sympathectomy and indomethacin injection, *Piezo2^fl/fl^; Ren^Cre^* mice and littermate controls were subjected to PEG-evoked hypovolemia or saline control treatment and blood plasma was isolated for measurement of renin (Fig. 5a). We found that plasma renin levels were strongly suppressed, in some cases to below the assay’s limit of detection, with blockade of both sympathetic and macula densa prostaglandin signaling in wild-type mice treated with saline (Fig. 5a), suggestive of a successful inhibition of renin-stimulating pathways. We also observed a substantial loss of TH+ sympathetic nerve fibers in the kidney, indicating the 6-OHDA treatment was efficacious (Extended Data Fig. 10a,b). Plasma renin was still elevated in response to PEG hypovolemia despite loss of these two important pathways, suggesting that a third (i.e. PIEZO2-dependent) pathway regulates renin levels after pharmacological blockade (Fig. 5b). Values observed in our wild-type littermate controls were lower than we had observed in our other hypovolemia experiments, indicating that the pharmacological manipulations did lower plasma renin levels (compare Fig. 5b to Fig. 4e). In *Piezo2^fl/fl^; Ren^Cre^* conditional knockout mice injected with saline, renin levels were largely unaffected by the blockade and similar to mice with intact sympathetic and macula densa signaling (compare to Fig. 2e). In PEG-injected conditional knockout mice, renin levels were substantially elevated beyond that of littermate controls and similar to hypovolemic *Piezo2^fl/fl^; Ren^Cre^* mice with intact sympathetic and macula densa signaling, suggesting that loss of mechanotransduction in renin-expressing cells dysregulates RAAS independently of the sympathetic and macular densa prostaglandin pathways (Fig. 5b). Although we cannot completely rule out remnant function of the macula densa after indomethacin treatment, the stark differences observed between the control and conditional knockout mice support our conclusions regardless of this possibility. Aldosterone levels were similarly exacerbated by hypovolemia in the knockouts, demonstrating that the changes in plasma renin translated into downstream RAAS signaling (Fig. 5c).

**Figure 5.**
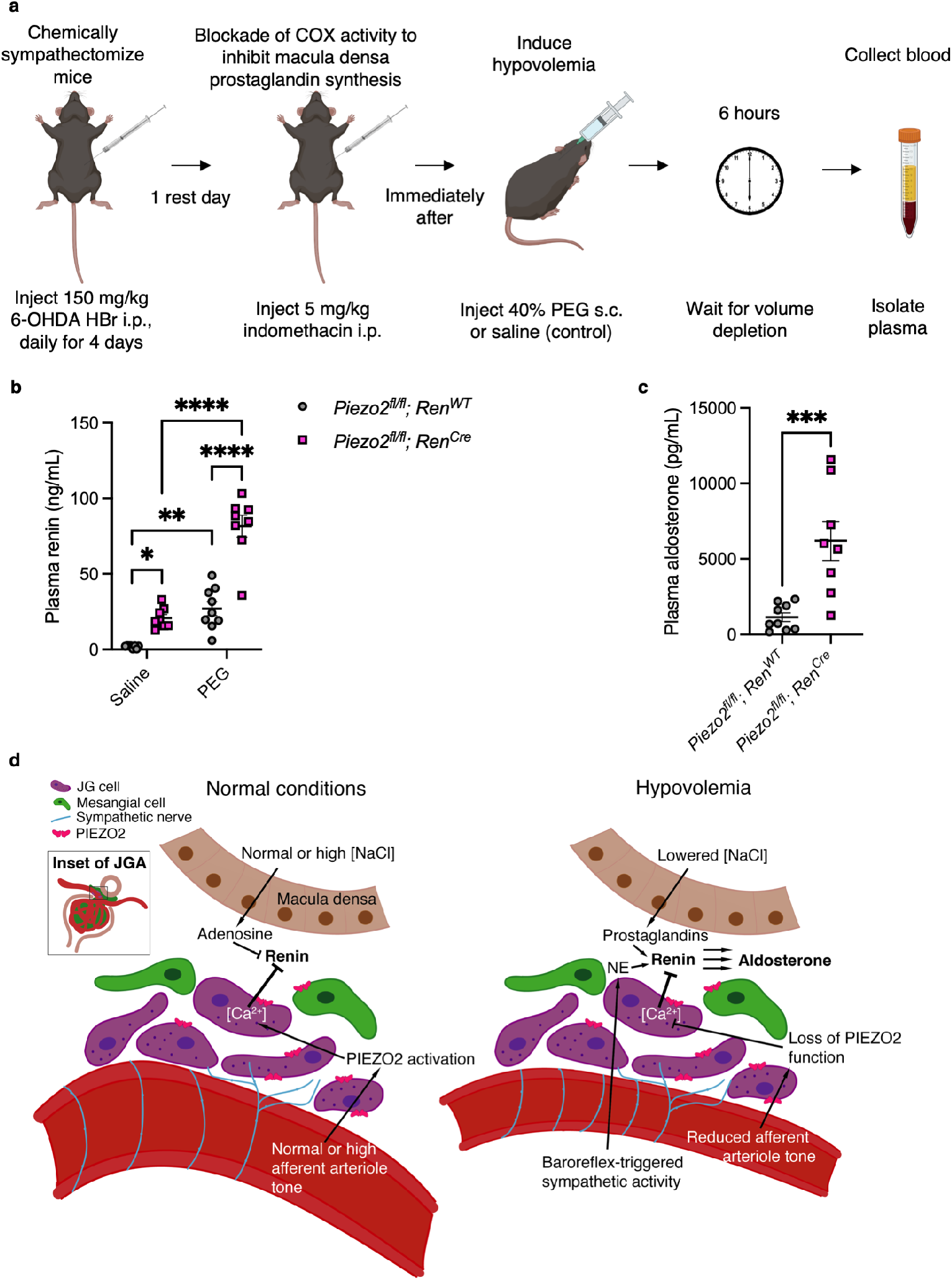
PIEZO2 contributes to RAAS independently of sympathetic and macula densa function. **a**, Experimental strategy to chemically ablate sympathetic efferent neurons and inhibit prostaglandin synthesis prior to induction of hypovolemia. **b**, Plasma renin levels in *Piezo2^fl/fl^; Ren^WT^* versus *Piezo2^fl/fl^; Ren^Cre^* animals after chemical sympathectomy and indomethacin treatment and six hours following saline or PEG (two-way ANOVA: \*\*\**p_interaction_* = 0.0004, F(1,29) = 15.81; Tukey’s multiple comparisons (left to right): * *p* = 0.0307, \*\**p* = 0.002, \*\*\*\**p* < 0.0001, \*\*\*\**p* < 0.0001; n = 8 *Ren^WT^* saline, 8 *Ren^Cre^* saline, 9 *Ren^WT^* PEG, and 8 *Ren^Cre^* saline). **c**, Plasma aldosterone levels in PEG-treated animals from **b** (Mann–Whitney: \*\*\**p* = 0.001, U = 4; n = 9 *Ren^WT^* and 8 *Ren^Cre^* mice). Each experiment was performed on at least two independent cohorts of mice, and error bars represent mean ± s.e.m. **d**, Schematic of model for role of PIEZO2 in regulation of renin in JG cells.

## Discussion

This work demonstrates that PIEZO2 in renin-expressing JG cells of the kidney regulates RAAS in response to changes in blood volume *in vivo* (Fig. 5d). Isolating the intrinsically mechanosensitive component of the renal baroreceptor is complicated due to the numerous intersecting feedback loops that encompass several organ systems, as well as the intrarenal chemical signaling pathways that modulate the activity of JG cells^64^. Our findings show that selectively perturbing the mechanosensory capability of renin-expressing cells through conditional deletion of *Piezo2* is sufficient to increase plasma renin in healthy mice. We demonstrate that elevated renin has downstream consequences for glomerular filtration rate, systemic blood pressure, and the hormonal response to hypovolemia. It can be challenging to reconcile the microscale changes in renal hemodynamics downstream of renin and TG feedback that were observed using micropuncture techniques in the rat^61,74,75^ with the whole-organism effects on RAAS we observed with selective gene perturbation in the mouse. Our findings suggest that altered JGA mechanotransduction has specific and profound outcomes for renal physiology *in vivo* and align with previous studies^61,74,75^. For example, we identified Ang(1-7)/Mas signaling as a major effector of PIEZO2 and renin’s effects on the GFR in the absence of elevated angiotensin II. Our data additionally support the conclusion that that PIEZO2 in renin-expressing cells acts independently of the parallel mechanisms (*i.e.* macula densa prostaglandin signaling and/or sympathetic activity) controlling renin levels during hypovolemia. Taken together, our study establishes a cellular and molecular mechanism for the response of the JGA to hypovolemia, underscores the therapeutic potential for targeting JG cell mechanotransduction pathways to modulate renal RAAS, and highlights the role of PIEZO proteins as general effectors of baroreception throughout the body.

## Methods

### Statistics

All statistical analyses were performed in Prism v9.5.0 (GraphPad). Error bars are defined as the mean ± s.e.m throughout, and individual data points are plotted. For blood pressure data in Figure 2, individual values separated by experimental subject are provided in the Extended Data Figures. All tested covariates are reported in the legends. Two-tailed tests were performed wherever applicable. *N*, test statistics, exact *p*-values and degrees of freedom (d.f.) are indicated where relevant in the figure legends. Normality and/or equal variance were not assumed and so nonparametric tests were used throughout.

### Study design

No analyses were performed in advance to pre-determine sample size. Sample sizes were based on similar studies in the literature^8,16,17^. No randomization was used. All *in vivo* experiments were performed by an experimenter blinded to the genotype of the animals tested. All experiments were independently repeated at least twice and data were pooled.

### Research animals

All experiments were approved by the Scripps Research Animal Care and Use Committee (Animal Use Protocol 08-0136). Mice were kept in standard housing with a 12-h light–dark cycle set with lights on from 6 am to 6 pm, with the room temperature kept around 22 °C, and humidity between 30% and 80% (not controlled). Mice were kept on pelleted paper bedding and provided with paper square nestlets and polyvinyl chloride pipe enrichment with *ad libitum* access to food and water. Age-matched littermate mice were used for all experiments. For all experiments, male and female mice were used and pooled. Mouse ages ranged from 12 to 20 weeks for all experiments unless otherwise indicated in the figure legends. All mice received metal identification tags (National Band & Tag, 1005-1) on the right ear when they were between 18 and 30 days old. After weaning between 21 and 30 days of age, mice were co-housed in groups of 2–5 littermates of the same sex. Genotyping was performed in-house by PCR from tail snip DNA samples using guidelines and primer sequences from Jackson Laboratory, or was performed by Transnetyx. The following strains of mice were used and maintained in the laboratory on an inbred background: *Piezo1^tdTomato^*(*B6;129-Piezo1^tm1.1Apat/^J*; Jackson Laboratories 029214) and *Piezo2^EGFP-IRES-Cre^* (*B6(SJL)-Piezo2^tm1.1(cre)Apat^/J*; Jackson Laboratories 027719). The following strains of mice were maintained on a *Ren1* monogenic C57BL6/J background: *Piezo1^fl/fl^* (*B6.Cg-Piezo1 ^tm2.1Apat^/J*; Jackson Laboratories 029213), *Piezo2^fl/fl^* (*B6(SJL)-Piezo2^tm2.2Apat^/J*, Jackson Laboratories 027720), *Ai9^fl/fl^* (*B6.Cg-Gt(ROSA)26Sor^tm9(CAG-tdTomato)Hze^/J*; Jackson Laboratories 007909), *Ai14^fl/fl^* (*B6.Cg-Gt(ROSA)26Sor^tm14(CAG-tdTomato)Hze^/J*; Jackson Laboratories 007914), *Pdgfrb^CreERT2^* (*B6.Cg-Pdgfrb^tm1.1(cre/ERT2)Csln^/J*, Jackson Laboratories 030201), *FoxD1^Cre^* (*B6;129S4-Foxd1^tm1(GFP/cre)Amc^/J*, Jackson Laboratories 012463), *Ren^Cre^* and *Ren^CreER^* (*Ren1c^Cre^* and *Ren1c^CreER^*, gifts from Drs. Kenneth Gross and Stuart Shankland^38,40^), and *SNS^Cre^*(*Tg(Scn10a-cre)^1Rkun^*, a gift from Dr. Rohini Kuner^56^, MGI: 3042874). Conditional knockout lines were maintained by crossing a homozygously floxed Cre-expressing mouse (homozygous for one or more indicated floxed alleles) with homozygously floxed mate. All strains are commercially available with the exception of *SNS^Cre^*, *Ren^Cre^*, and *Ren^CreER^*. For experiments involving the *Pdgfrb^CreERT2^*line, recombination was achieved with once-daily intraperitoneal injection of 75 mg per kg body weight tamoxifen (Sigma-Aldrich, T5648) dissolved in 0.22-µm sterile-filtered corn oil delivered to both Cre-expressing and control mice over five consecutive days when between 8-12 weeks of age. Mice were used four or more weeks after tamoxifen administration to ensure adequate time for Cre activity and protein turnover. In all experiments with the *Pdgfrb^CreERT2^*line, both littermate controls and Cre-expressing animals received tamoxifen.

### Single molecule fluorescent in situ hybridization (smFISH)

For mouse experiments, kidneys were removed immediately, embedded in optimal cutting temperature compound (OCT, Sakura), and flash-frozen in liquid nitrogen. For human kidney biopsies, 5 µm-thickness formalin-fixed paraffin embedded (FFPE) kidney sections were obtained from the Kidney Translational Resource Center at Washington University and processed using the manufacturer’s instructions for FFPE slides. Tissue from a single anonymous White/Caucasian male donor aged 46 was used. Informed consent and IRB approval for human kidney samples was obtained by the KTRC. The protocol for RNAscope Multiplex Fluorescent Reagent Kit V2 (ACDBio, 323100) was followed exactly according to the instructions for fresh-frozen and FFPE tissue. Protease IV was applied for 30 min for mouse tissue when only smFISH was performed. When IHC was performed following smFISH, Protease III was applied for 30 minutes instead and the manufacturer’s instructions for IHC following smFISH were followed exactly, using a Rabbit anti-NPHS2 primary antibody (Abcam, ab50339) at 1:1000 followed by goat anti-Rabbit HRP-conjugated secondary antibody (Abcam, ab6721) at 1:1000. For human kidney sections, manufacturer’s instructions were followed exactly for FFPE kidney tissue. Probes (all from ACDBio) for mouse *Piezo1* (#400181), mouse *Piezo2* (#400191), mouse *Piezo2*-E43-E45 (#439971), mouse *Ren1* (#433461), mouse *Pdgfrb* (#411381), mouse *Pecam1* (#316721), human *PIEZO1* (#485101), human *PIEZO2* (#449951), and human *PDGFRB* (#548991) were applied to detect transcript. The manufacturer’s 3-plex negative control probe (#320871) was used in each experiment to detect non-specific signal. Displayed images were uniformly cropped from the original images.

### Immunohistochemistry (IHC)

For *Piezo1^tdTomato^* and *Piezo2^GFP-Cre^* IHC experiments, tissues were processed using a modified protocol to preserve signal^35^. In brief, fresh-frozen kidneys were embedded in OCT and sectioned at 20 µm. Sections were post-fixed on slides in cold 4% paraformaldehyde (PFA) in PBS for 10 min at room temperature and quenched using 20 mM glycine and 75 mM ammonium chloride with 0.1% v/v Triton X-100 in PBS for 10 min. Slides were washed in PBS and then incubated in blocking buffer (0.6% w/v fish skin gelatin with 0.05% w/v saponin in PBS with 5% v/v normal goat or donkey serum) for 1 h at room temperature. Slides were incubated in primary antibodies overnight at 4 °C in blocking buffer without serum: AlexaFluor 647-conjugated FluoTag-X4 anti-RFP single domain antibody (Nanotag, N0404, 1:100) or chicken anti-GFP (Aves Labs, GFP1010, 1:1000). When conjugated nanobody was used, slides were washed in PBS and mounted in SlowFade Diamond immediately prior to imaging. For GFP staining experiments, slides were washed in PBS, and then incubated in goat anti-chicken Alexa Fluor 488 secondary antibody (Life Technologies, A11039, 1:1000) in blocking buffer 1 h at room temperature. Samples were washed in PBS, counterstained with 1:30,000 TO-PRO-3 Iodide (Life Technologies, T3605), and then mounted in SlowFade Diamond mounting medium (Life Technologies, S36967) and sealed with nail polish prior to imaging. For conventional IHC, fresh-frozen kidneys were embedded in OCT and sectioned at 20 µm. Sections were post-fixed on slides for 15 minutes at 4 °C in 4% v/v PFA-PBS, briefly rinsed in PBS, washed for 10 min in 0.3% v/v Triton X-100 in PBS (PBST), then blocked for 1 h in 5% v/v normal goat serum in 0.3% PBST. Sections were incubated overnight at 4 °C in rabbit anti-renin (Abcam, ab212197, 1:250), rat anti-PECAM1 (Sigma Aldrich, CBL-1337-1, 1:1000), rabbit anti-NPHS2 (Abcam, ab50339, 1:1000), or rabbit anti-RFP (Rockland, 600-401-379, 1:1000) in 0.3% PBST with 1% NGS. Sections were washed in PBS and incubated in 1:1,000 goat anti-rabbit AlexaFluor 647 (Life Technologies, A21245) and/or goat anti-rat 488 (Life Technologies, A11006) for 1 h at room temperature. Tissues were rinsed in PBS, mounted in HIGHDEF IHC Fluoromount (Enzo), and sealed with nail polish. All images were acquired on either a Nikon A1 (20x air objective) or AX (16x water immersion objective) confocal microscope and the imaging settings (laser power, gain, 1,024 × 1,024 original resolution, pixel dwell, objective and use of Nyquist zoom) were kept consistent within experiments. For all images, brightness and contrast adjustments were uniformly applied to the entire image. Images were processed and analyzed using FIJI (ImageJ2 v2.3.0/1.53f).

### Blood collection methods

For terminal experiments, mice were euthanized via isoflurane overdose and 0.3-0.7 mL whole blood was collected after decapitation. For non-terminal experiments, mice were anesthetized with 3% isoflurane/2% oxygen and < 200 µL whole blood was collected from the retroorbital sinus using a sterile micropipette tip with a fire-polished end. All blood samples were collected into lithium heparin-coated tubes (BD Microtainer #365965). Blood was spun at 1200g for 20 min immediately following collection. Plasma supernatants were collected and stored at −80 °C for up to a month prior to assay, and only freeze-thawed once. Blood was collected between 2-5pm.

### Enzyme-linked immunosorbent assays (ELISA)

ELISA was performed using the indicated assays for the following analytes according to the manufacturer’s instructions: renin (LSBio, LS-F508-1), aldosterone (Tecan, RE52301), angiotensin II (Ray Biotech, EIA-ANGII-1), albumin (Abcam, ab108792). The appropriate plasma or urine dilution was empirically determined using a dilution series. Standard curves and extrapolation of sample concentrations were determined using a 4-parameter logistic fit in Prism 9.5.0. Standards and negative controls were run on each plate, and all samples and standards were run in duplicate for each assay. Plates were read according to manufacturer’s instructions using the Cytation 3 plate reader (Agilent) with Gen5 software (v2.04).

### Volume pressure recording measurement of systemic blood pressure

The CODA High Throughput VPR System (Kent Scientific) was used for all experiments according to published methods^45^. Briefly, mice were habituated to the appropriately sized rodent restrainer, cuff set, and heated platform for 5 days prior to measurements. The same restrainer was used for each mouse for the duration of the experiment (wiping only with deionized water and lint-free tissue) to habituate the mouse to familiar odors and was stored in a sealed plastic bag in between habituations. Only cage mates were tested in parallel to reduce stress. Tail temperature was verified with an infrared thermometer prior to beginning measurements and mice were tested when tail temperature was between 32-35 °C. For each day of measurements (3 per mouse), 10 acclimation and 20 experimental cycles were performed. Only measurements that passed software quality control (CODA Data Acquisition Software, version 1.06) were analyzed.

### Non-invasive transdermal measurement of glomerular filtration rate (GFR)

GFR was measured using the transdermal system from MediBeacon^49,53–55^. Briefly, mouse flanks were dehaired with depilatory cream (Nair) on the day prior to measurement. Mice were anesthetized with 2% isoflurane/1% oxygen and placed on a 37 °C heating pad. The transducer was applied to the dehaired flank skin of the mouse using the supplied adhesive patch, avoiding any pigmented skin regions. Baseline was acquired for 1-3 min. Mice received a bolus injection of 2.5 µL/g b.w. of 30 mg/mL fluorescein isothiocyanate (FITC)-sinistrin (MediBeacon) prepared in sterile PBS and delivered via the retroorbital sinus with a 28G insulin syringe. Measurements were acquired for one hour post-injection and data were analyzed and fitted offline using the MBLab2 software (MediBeacon, v2.12) according to the manufacturer’s instruction to calculate the *t_1/2_* in minutes. GFR was calculated from *t_1/2_* using the formula for adult C57BL6/J mice^49^:

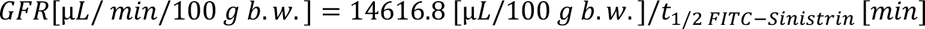

### Urine collection methods

Mice were lightly scruffed one at a time over sterile hydrophobic LabSand (Braintree Scientific) until they urinated and for no more than 30 sec. This method reliably yielded between 10-100 µL urine per mouse. Urine was immediately collected using a clean micropipette tip and centrifuged at 800g for 10 min at 4 °C. Urine was stored at −80 °C until assessment via ELISA. Urine was harvested at 2pm.

### Histology

Hematoxylin and eosin (H&E) and Periodic-acid Schiff (PAS) staining were performed on formalin-fixed mouse kidney sections by the Sanford Burnham Prebys Histology Core (La Jolla, CA) using standard paraffin embedding, sectioning, and staining methods. The slides were analyzed for glomerulopathy, mesangial cell number, mesangial matrix deposition, tubular morphology, and quantity of nuclei in the juxtaglomerular apparatus by an individual blinded to genotypes. In total, sections from n = 4 *Piezo2^fl/fl^; Pdgfrb^WT^* ; n = 4 *Piezo2^fl/fl^; Pdgfrb^CreERT2^* n = 7 *Piezo2^fl/fl^; Ren^WT^*, and n = 4 *Piezo2^fl/fl^; Ren^Cre^* mice were analyzed. Representative images of kidney cortex were acquired using a Keyence BZ-X710 microscope using brightfield imaging with a 40x objective and the supplied color camera.

### Captopril administration

Water valves were removed from cages and mice were supplied with 400mg/L captopril (Sigma-Aldrich, C4042) in the drinking water, prepared fresh each day in a water bottle. Mice had access to standard chow. After six days of treatment, GFR was measured. On the seventh day, blood was harvested for ELISA.

### A779 administration

Mice were injected i.p. once daily for seven consecutive days with 0.5 mg/kg A779 (Cayman Chemical, 23396) dissolved in 0.9% NaCl in water.

### Polyethylene glycol (PEG) mouse model of hypovolemia

Mice were lightly and briefly (< 5 min) anesthetized with 2% isoflurane and injected with 40% w/v PEG-8000 (Sigma-Aldrich, 89510) in sterile 0.9% NaCl subcutaneously using a 28G insulin syringe. Food and water were removed from the cage. After six hours, mice were euthanized and blood was collected. For the 6-OHDA and indomethacin experiments, 15 minutes after indomethacin injection, mice received either sterile saline or 40% PEG. Blood was collected six hours following PEG or saline injection.

### 6-OHDA and indomethacin administration

Mice were injected i.p. once daily for four consecutive days with 150 mg/kg 6-OHDA HBr (Sigma-Aldrich, 162957) dissolved in 0.02% ascorbic acid and 0.9% NaCl in water. 24 hours after the final injection, mice were injected i.p. with 5 mg/kg indomethacin (Tocris, 1708) dissolved in 0.01 M sodium carbonate with 1% DMSO in water.

## Data availability

All data points are presented as dot plots in the Figures or Extended Data Figures. Raw data are available upon reasonable request from the authors.

## Code availability

N/A

## Acknowledgements

The authors thank all members of the Patapoutian laboratory for providing helpful feedback, A. Dubin (Scripps Research) for critical editing of the manuscript, G. Garcia (Sanford Burnham Prebys Medical Discovery Institute) for histology services, the Dorris Neuroscience Center Department of Animal Resources for animal husbandry services, K. Conlon and S. Jain at the Kidney Translational Resource Center (supported by the Division of Nephrology at Washington University) for processing and providing human kidney samples and obtaining IRB approval and informed consent, V. Augustine (University of California San Diego) for advice on the hypovolemia assay, K. Spencer and the Nikon Center of Excellence Imaging Center (Scripps Research) for imaging facilities, S. Shankland (University of Washington) and K. Gross (Roswell Park Comprehensive Cancer Center) for the gift of the *Ren^Cre^*mice, and R. Kuner (Heidelberg University) for the gift of the *SNS^Cre^*mice.

## Contributions

Conceptualization: RZH, AP

Methodology: RZH, JHM, SB

Investigation: RZH, SS, SB, JHM

Visualization: RZH

Funding acquisition: RZH, AP

Project administration: RZH, AP

Supervision: RZH, AP

Writing – original draft: RZH

Writing – review & editing: RZH, JHM, SS, SB, AP

## Supplementary information

No supplementary information is included.

## Corresponding authors

Correspondence to Ardem Patapoutian or Rose Z. Hill.

## Ethics declarations

*Competing interests:* The authors declare no competing interests.

## Extended data figure legends

**Extended Data Figure 1.**
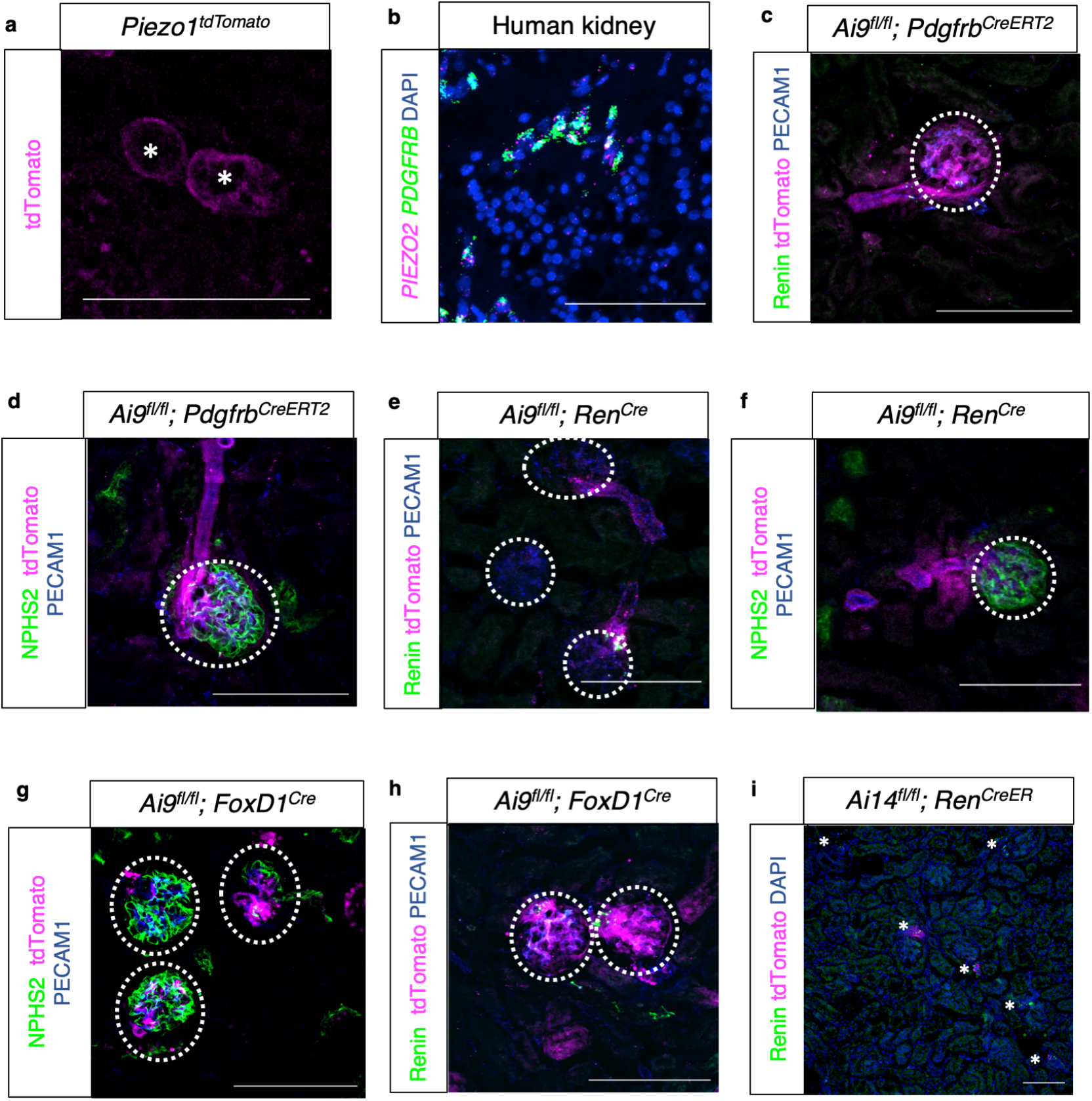
**a**, Sectioned mouse kidney stained with anti-tdTomato AlexaFluor 647-conjugated nanobody. Asterisk (*) indicates distal convoluted tubule. **b**, smFISH of sectioned human kidney for *PIEZO2*, *PDGFRB*, and counterstained with DAPI. **c**, Sectioned mouse kidney with native tdTomato fluorescence, stained with anti-Renin and anti-PECAM1 antibodies. **d**, Sectioned mouse kidney with native tdTomato fluorescence, stained with anti-NPHS2 and anti-PECAM1 antibodies. **e**, Sectioned mouse kidney with native tdTomato fluorescence, stained with anti-Renin and anti-PECAM1 antibodies. **f**, Sectioned mouse kidney with native tdTomato fluorescence, stained with anti-NPHS2 and anti-PECAM1 antibodies. **g**, Sectioned mouse kidney with native tdTomato fluorescence, stained with anti-NPHS2 and anti-PECAM1 antibodies. **h**, Sectioned mouse kidney with native tdTomato fluorescence, stained with anti-Renin and anti-PECAM1 antibodies. Dotted circles outline renal corpuscles. **i**, Sectioned mouse kidney with native tdTomato fluorescence, stained with anti-Renin antibodies. Asterisks (*) are placed to the immediate left of JGA. Scale bars = 100 µm. Each experiment was repeated on N=2 mice.

**Extended Data Figure 2.**
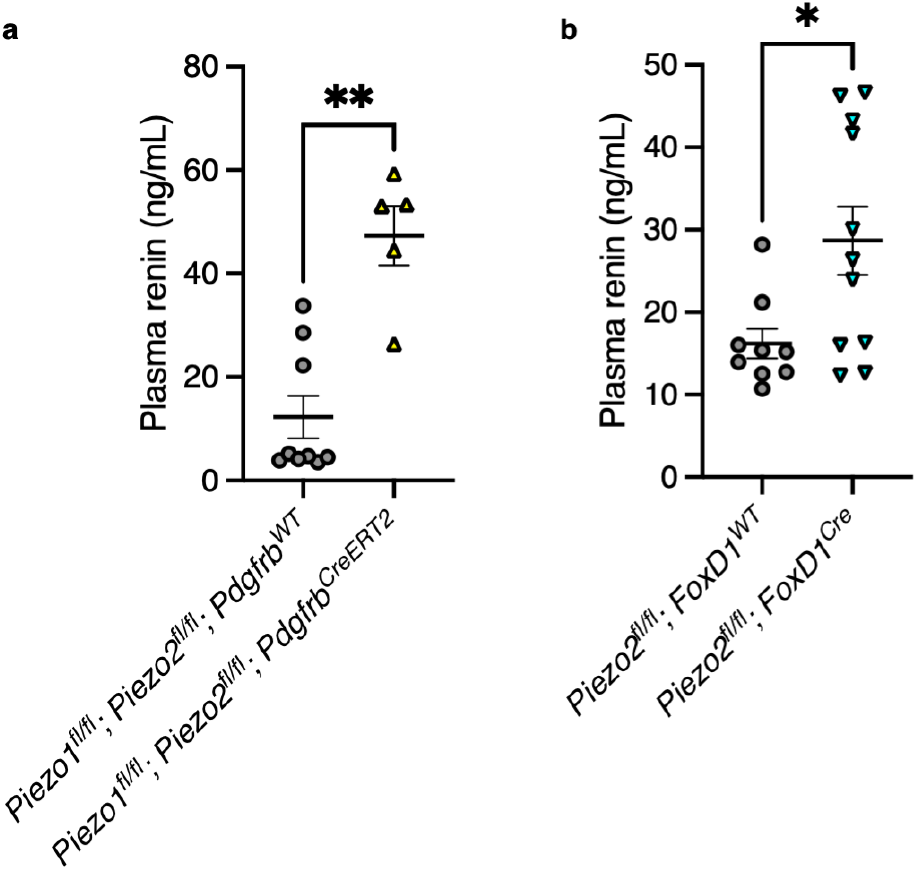
**a**, Plasma renin levels in *Piezo1^fl/fl^; Piezo2^fl/fl^; Pdgfrb^WT^* versus *Piezo1^fl/fl^; Piezo2^fl/fl^; Pdgfrb^CreERT2^* animals (Mann–Whitney: \*\**p* = 0.0040, U = 2; n = 9 *Pdgfrb^WT^* and 5 *Pdgfrb^CreERT2^* mice). **b**, Plasma renin levels in *Piezo2^fl/fl^; FoxD1^WT^* versus *Piezo2^fl/fl^; FoxD1^Cre^* animals (Mann–Whitney: \**p* = 0.0381, U = 22; n = 9 *FoxD1^WT^* and 11 *FoxD1^Cre^* mice). Each experiment was performed on at least two independent cohorts of mice, and error bars represent mean ± s.e.m.

**Extended Data Figure 3.**
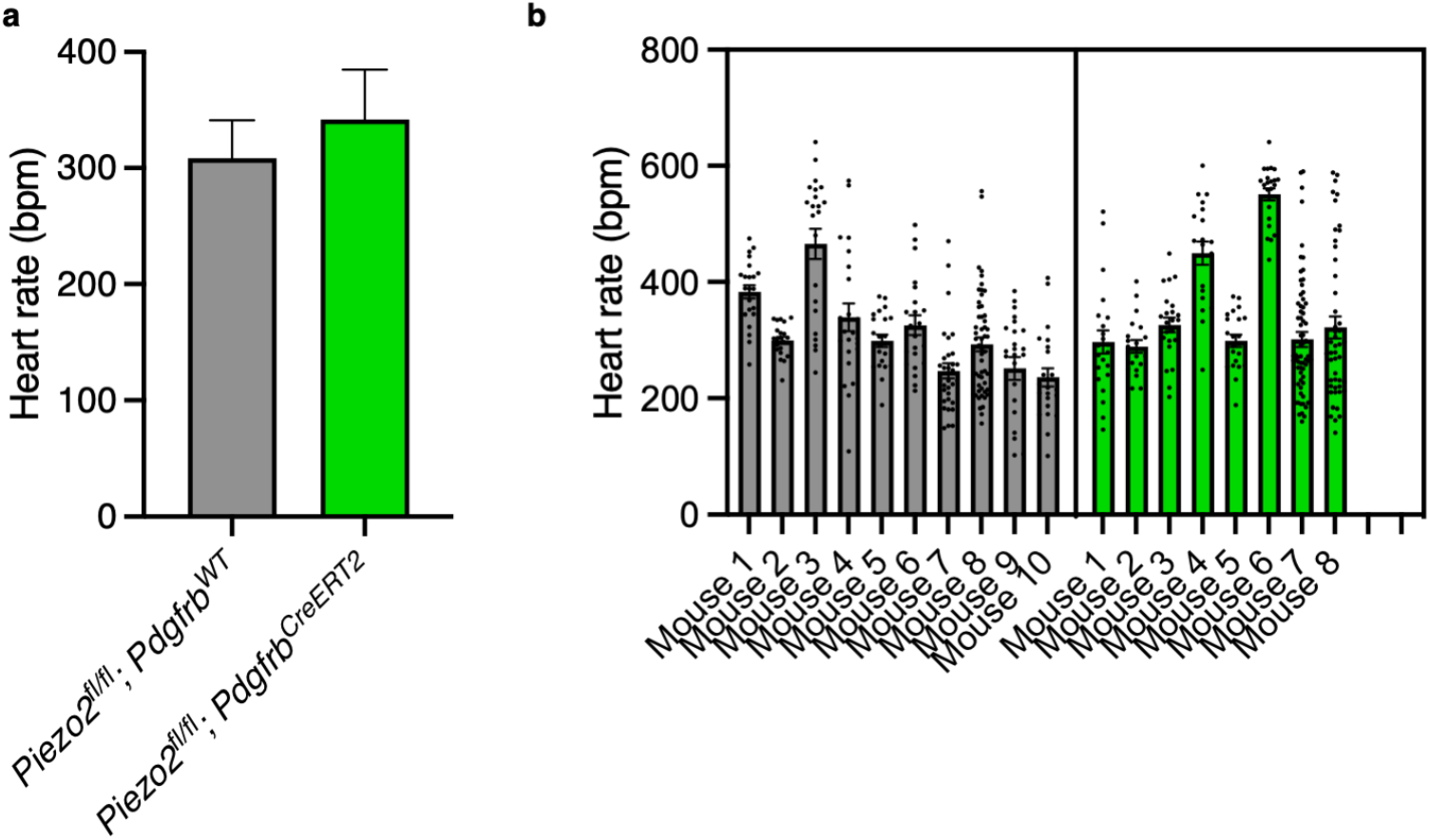
**a**, Heart rate (beats per minute) measured using the VPR system in *Piezo2^fl/fl^; Pdgfrb^WT^* versus *Piezo2^fl/fl^; Pdgfrb^CreERT2^* animals (two-tailed nested *t*-test: = 0.3122, *t* = 1.044, d.f. = 16, n = 10 *Pdgfrb^WT^* and 8 *Pdgfrb^CreERT2^* mice). **b**, Data in **a** replotted to show individual data points per mouse, with *Piezo2^fl/fl^; Pdgfrb^WT^* in gray and *Piezo2^fl/fl^; Pdgfrb^CreERT2^* in green. Experiment was performed on two independent cohorts of mice, and error bars represent mean ± s.e.m.

**Extended Data Figure 4.**
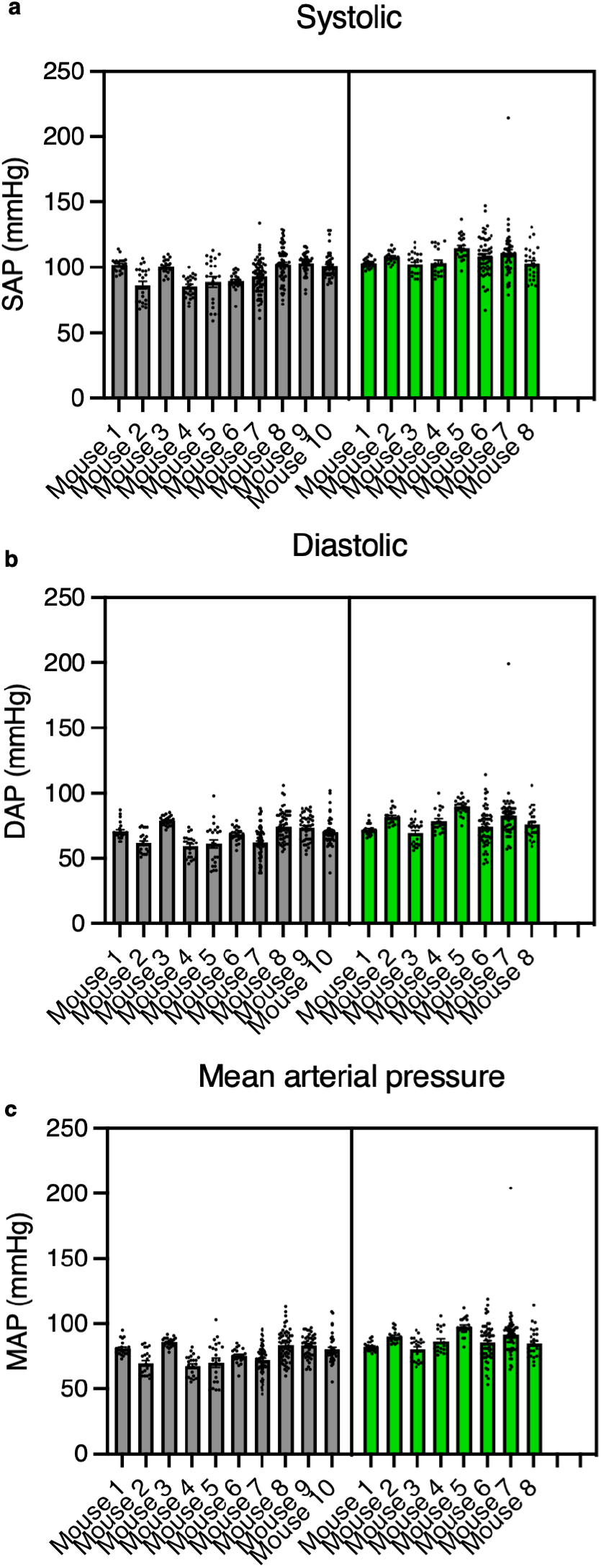
**a**, Systolic blood pressure data from *Piezo2^fl/fl^; Pdgfrb^WT^* (gray) versus *Piezo2^fl/fl^; Pdgfrb^CreERT2^*(green) animals replotted from Fig. 2d to show all trials from individual mice. **b**, Diastolic blood pressure data from *Piezo2^fl/fl^; Pdgfrb^WT^* (gray) versus *Piezo2^fl/fl^; Pdgfrb^CreERT2^* (green) animals replotted from Fig. 2d to show all trials from individual mice. **c**, Mean arterial blood pressure data from *Piezo2^fl/fl^; Pdgfrb^WT^* (gray) versus *Piezo2^fl/fl^; Pdgfrb^CreERT2^* (green) animals replotted from Fig. 2d to show all trials from individual mice. Experiment was performed on two independent cohorts of mice, and error bars represent mean ± s.e.m.

**Extended Data Figure 5.**
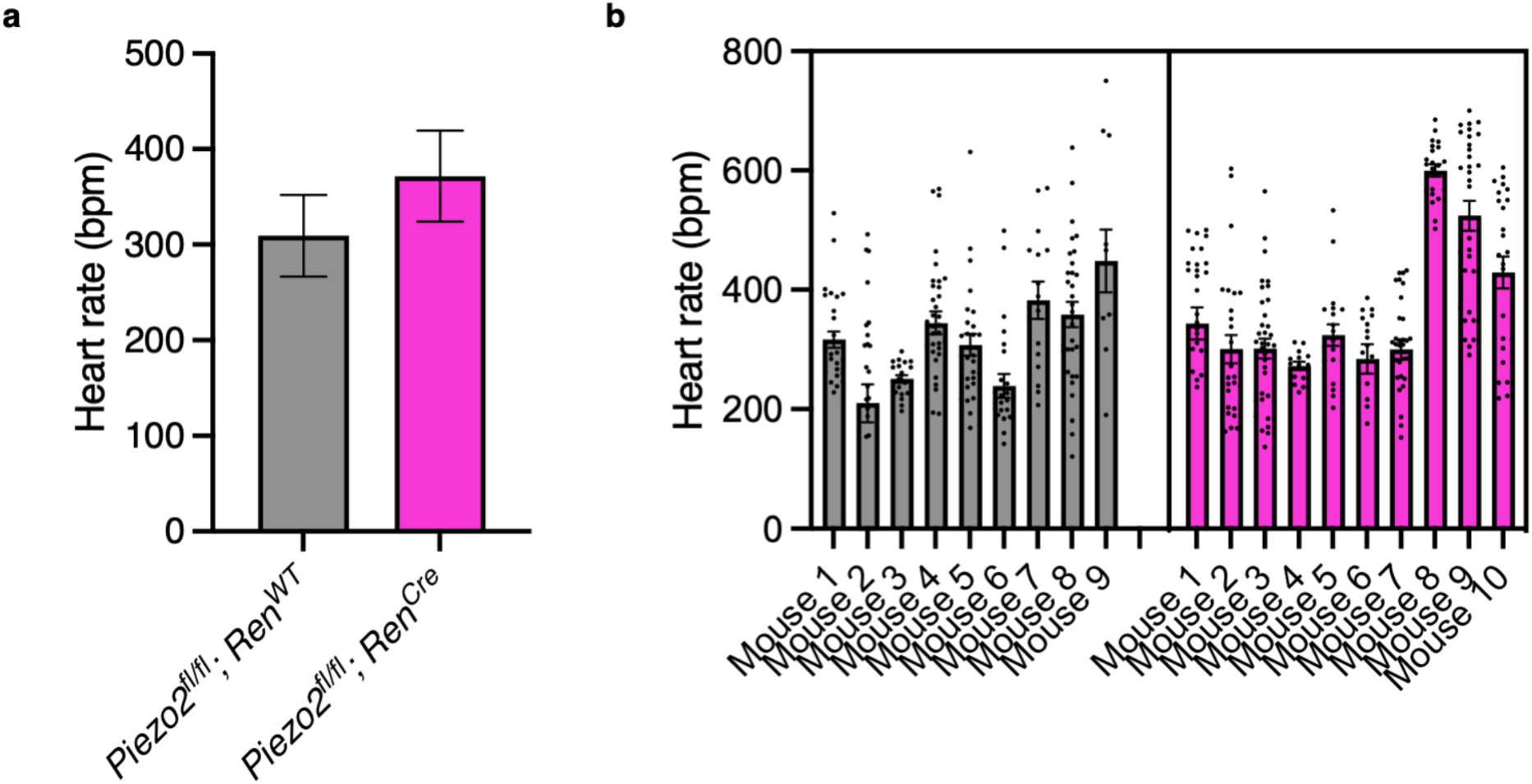
**a**, Heart rate (beats per minute) measured using the VPR system in *Piezo2^fl/fl^; Ren^WT^* versus *Piezo2^fl/fl^; Ren^Cre^* animals (two-tailed nested *t*-test: = 0.2572, *t* = 1.374, d.f. = 17, n = 10 *Ren^WT^* and 9 *Ren^Cre^* mice). **b**, Data in **a** replotted to show individual data points per mouse, with *Piezo2^fl/fl^; Ren^WT^* in gray and *Piezo2^fl/fl^; Ren^Cre^* in magenta. Experiment was performed on two independent cohorts of mice, and error bars represent mean ± s.e.m.

**Extended Data Figure 6.**
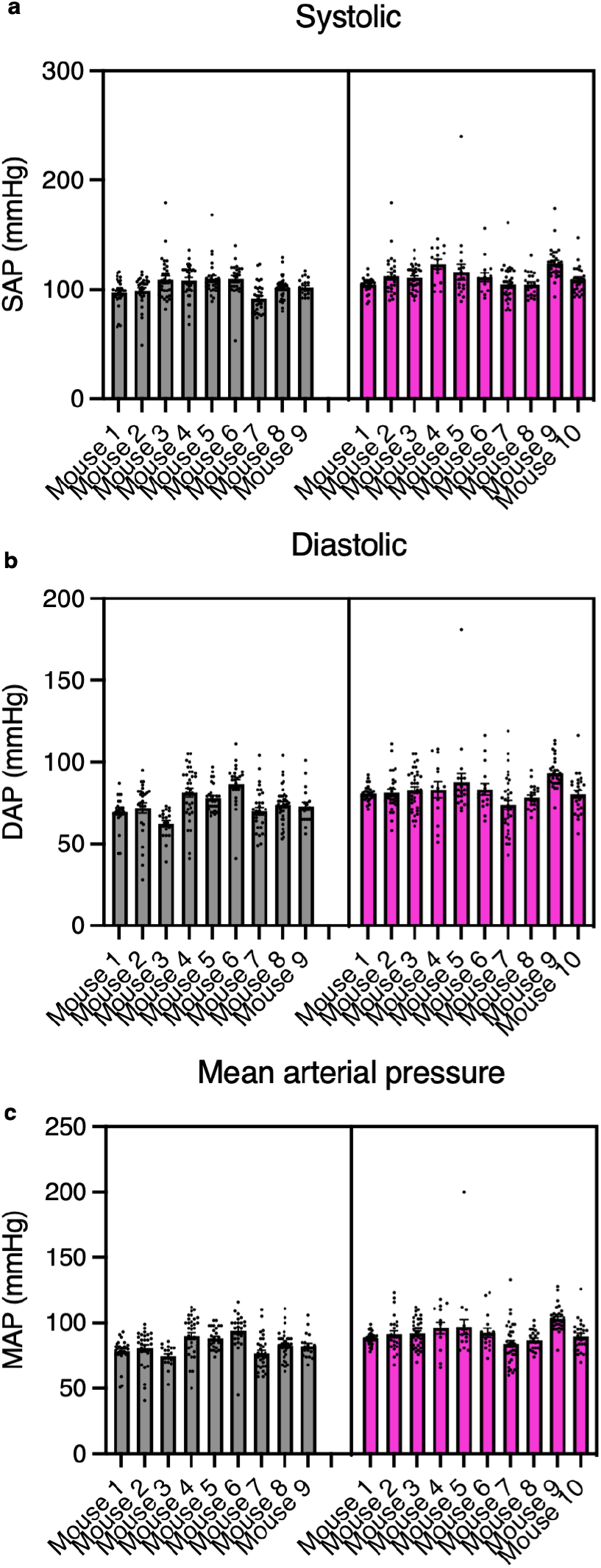
**a**, Systolic blood pressure data from *Piezo2^fl/fl^; Ren^WT^* (gray) versus *Piezo2^fl/fl^; Ren^Cre^* (magenta) animals replotted from Fig. 2f to show all trials from individual mice. **b**, Diastolic blood pressure data from *Piezo2^fl/fl^; Ren^WT^* (gray) versus *Piezo2^fl/fl^; Ren^Cre^* (magenta) animals replotted from Fig. 2f to show all trials from individual mice. **c**, Mean arterial blood pressure data from *Piezo2^fl/fl^; Ren^WT^* (gray) versus *Piezo2^fl/fl^; Ren^Cre^* (magenta) animals replotted from Fig. 2f to show all trials from individual mice. Experiment was performed on two independent cohorts of mice, and error bars represent mean ± s.e.m.

**Extended Data Figure 7.**
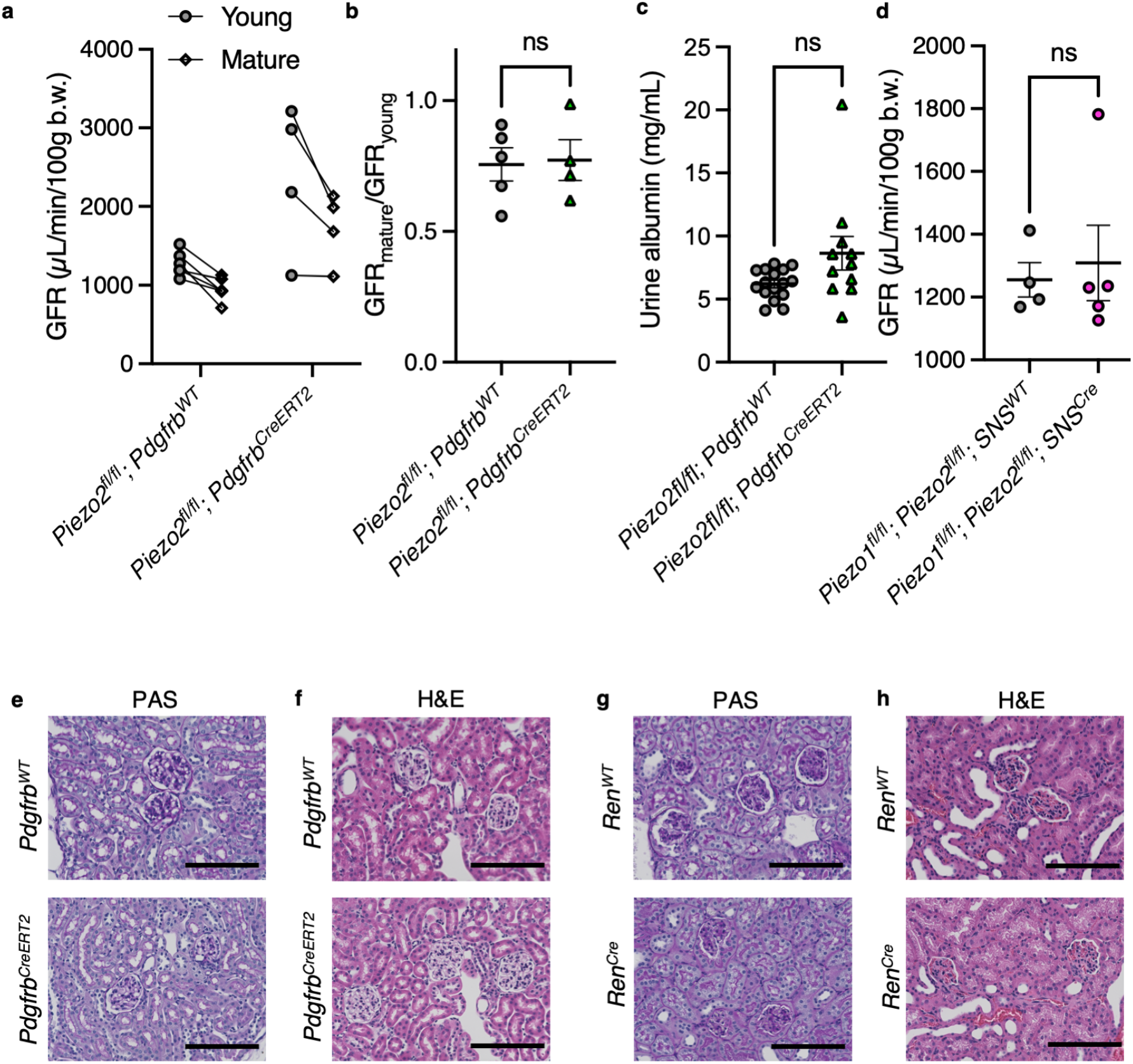
**a**, GFR measured in young (3-6 months) and then mature (11-14 months) *Piezo2^fl/fl^; Pdgfrb^WT^*and *Piezo2^fl/fl^; Pdgfrb^CreERT2^* animals. **b**, Comparison of the ratio of (GFR_mature_/GFR_young_) plotted in **a** (Mann–Whitney: *p* > 0.9999, U = 10; n = 5 *Pdgfrb^WT^* and 4 *Pdgfrb^CreERT2^* mice). **c**, Albumin concentration measured from urine of *Piezo2^fl/fl^; Pdgfrb^WT^* and *Piezo2^fl/fl^; Pdgfrb^CreERT2^* animals (Mann–Whitney: *p* = 0.0547, U = 49; n = 16 *Pdgfrb^WT^* and 11 *Pdgfrb^CreERT2^* mice). **d**, GFR in *Piezo1^fl/fl^; Piezo2^fl/fl^; SNS^WT^* versus *Piezo1^fl/fl^; Piezo2^fl/fl^; SNS^Cre^* animals (Mann–Whitney: *p* = 0.9048, U = 9; n = 4 *SNS^WT^* and 5 *SNS^Cre^* mice). Each experiment was performed on at least two independent cohorts of mice, and error bars represent mean ± s.e.m. **e**, PAS staining of *Piezo2^fl/fl^; Pdgfrb^WT^* (upper) and *Piezo2^fl/fl^; Pdgfrb^CreERT2^* (lower) kidney sections. **f**, H&E staining of *Piezo2^fl/fl^; Pdgfrb^WT^* (upper) and *Piezo2^fl/fl^; Pdgfrb^CreERT2^* (lower) kidney sections. **g**, PAS staining of *Piezo2^fl/fl^; Ren^WT^* (upper) and *Piezo2^fl/fl^; Ren^Cre^* (lower) kidney sections. **h**, H&E staining of *Piezo2^fl/fl^; Ren^WT^* (upper) and *Piezo2^fl/fl^; Ren^Cre^* (lower) kidney sections. Images are representative of n = 4 *Piezo2^fl/fl^; Pdgfrb^WT^* ; n = 4 *Piezo2^fl/fl^; Pdgfrb^CreERT2^* n = 7 *Piezo2^fl/fl^; Ren^WT^*, and n = 4 *Piezo2^fl/fl^; Ren^Cre^* mice (see Methods). Scale bars = 100 µm.

**Extended Data Figure 8.**
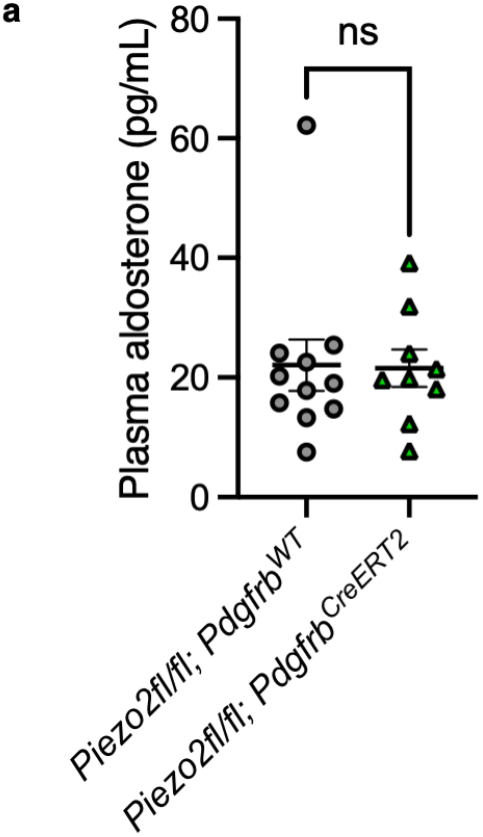
**a**, Plasma aldosterone levels in *Piezo2^fl/fl^; Pdgfrb^WT^* versus *Piezo2^fl/fl^; Pdgfrb^CreERT2^* animals after seven days of captopril (Mann–Whitney: *p* = 0.7664, U = 45; n = 11 *Pdgfrb^WT^* and 9 *Pdgfrb^CreERT2^*mice). Experiment was performed on at least two independent cohorts of mice, and error bars represent mean ± s.e.m.

**Extended Data Figure 9.**
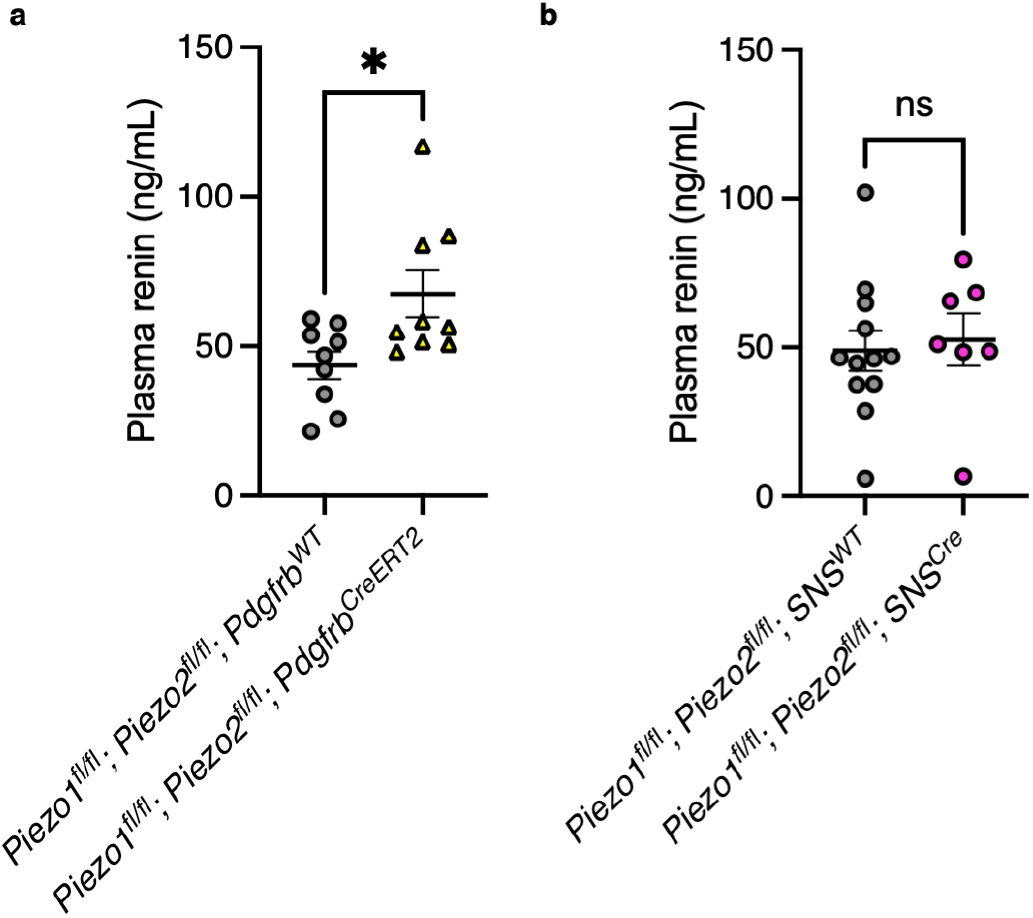
**a**, Plasma renin levels in *Piezo1^fl/fl^; Piezo2^fl/fl^; Pdgfrb^WT^* versus *Piezo1^fl/fl^; Piezo2^fl/fl^; Pdgfrb^CreERT2^* animals six hours following PEG injection (Mann–Whitney: \**p* = 0.0315, U = 16; n = 9 *Pdgfrb^WT^* and 9 *Pdgfrb^CreERT2^* mice). **b**, Plasma renin levels in *Piezo1^fl/fl^; Piezo2^fl/fl^; SNS^WT^* versus *Piezo1^fl/fl^; Piezo2^fl/fl^; SNS^Cre^* animals six hours following PEG injection (Mann–Whitney: *p* = 0.2614, U = 28; n = 12 *SNS^WT^* and 7 *SNS^Cre^* mice). Each experiment was performed on at least two independent cohorts of mice, and error bars represent mean ± s.e.m.

**Extended Data Figure 10.**
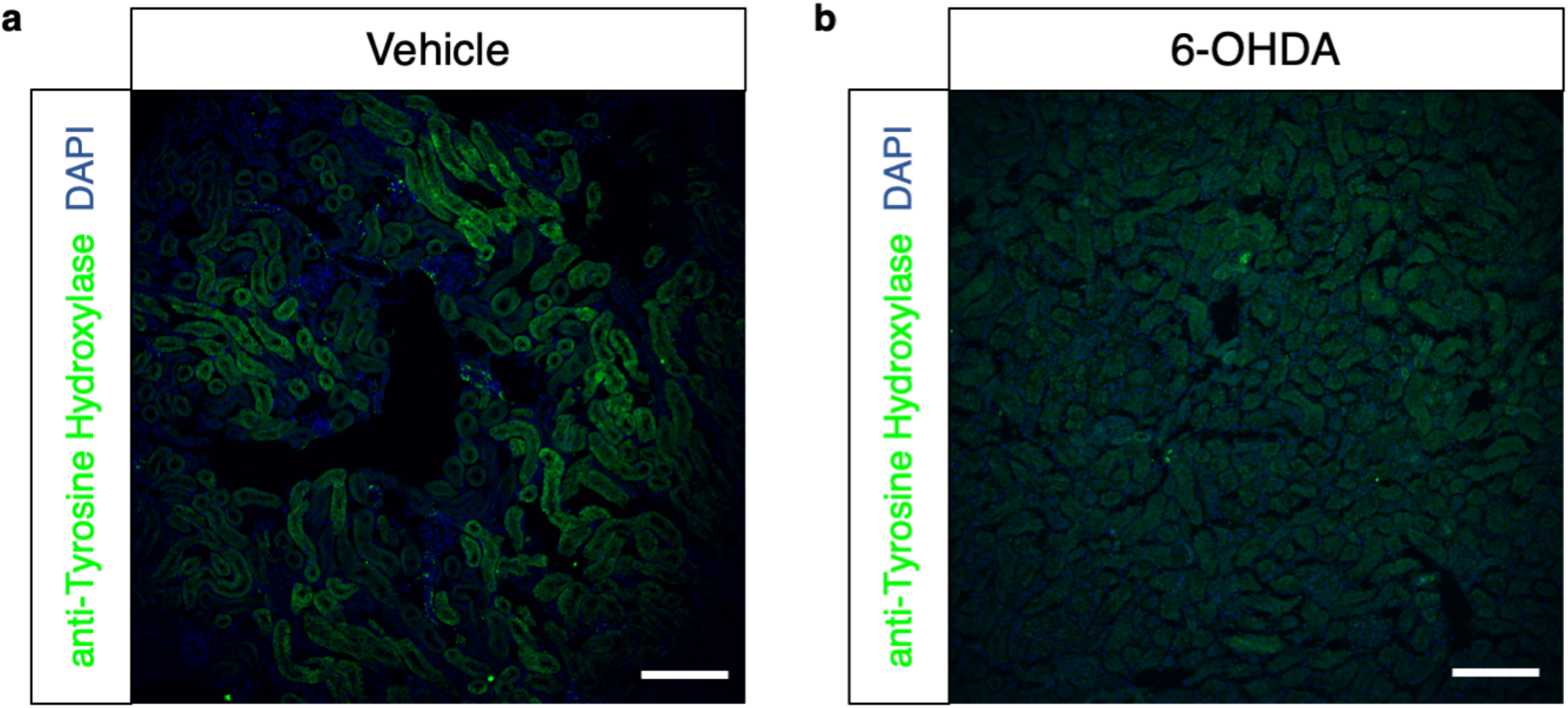
**a**, Sectioned mouse kidney stained with anti-tyrosine hydroxylase antibody and DAPI after vehicle treatment. **b**, Sectioned mouse kidney stained with anti-tyrosine hydroxylase antibody and DAPI after 6-OHDA treatment. Scale bars = 100 µm. Experiment was performed on N=2 mice.

